# Deciphering the many maps of the Xingu – an assessment of land cover classifications at multiple scales

**DOI:** 10.1101/2019.12.23.887588

**Authors:** M Kalacska, J.P Arroyo-Mora, O Lucanus, L Sousa, T Pereira, T Vieira

## Abstract

Remote sensing is an invaluable tool to objectively illustrate the rapid decline in habitat extents worldwide. The many operational Earth Observation platforms provide options for the generation of land cover maps, each with unique characteristics, as well as considerable semantic differences in the definition of classes. As a result, differences in baseline estimates are inevitable. Here we compare forest cover and surface water estimates over four time periods spanning three decades (1989–2018) for ∼1.3 million km^2^ encompassing the Xingu river basin, Brazil, from published, freely accessible remotely sensed classifications. While all datasets showed a decrease in forest extent over time, we found a large range in the total area reported by each product for all time periods. The greatest differences ranged from 9% (year 2000) to 17% of the total area (2014-2018 period). We also show the high sensitivity of forest fragmentation metrics (entropy and foreground area density) to data quality and spatial resolution, with cloud cover and sensor artefacts resulting in errors. We further show the importance of choosing surface water datasets carefully because they differ greatly in location and amount of surface water mapped between sources. In several of the datasets illustrating the land cover following operationalization of the Belo Monte dam, the large reservoirs are notably absent. Freshwater ecosystem health is influenced by the land cover surrounding water bodies (e.g. Riparian zones). Understanding differences between the many remotely sensed baselines is fundamentally important to avoid information misuse, and to objectively choose the most appropriate dataset for conservation, taxonomy or policy-making. The differences in forest cover between the datasets examined here are not a failure of the technology, but due to different interpretations of ‘forest’ and characteristics of the input data (e.g. spatial resolution). Our findings demonstrate the importance of transparency in the generation of remotely sensed datasets and the need for users to familiarize themselves with the characteristics and limitations of each chosen data set.

## Introduction

Earth observation from satellite imagery is fundamentally important for conservation (Rose et al. 2015; Li et al. 2016). Since the early days of deforestation mapping from satellites (e.g. Townshend and Justice 1988; Westman et al. 1989; Skole and Tucker 1993), remote sensing has become an invaluable tool for objectively illustrating the rapid decline in habitat extent (Pimm et al. 2001; O’Connor et al. 2015; Paganini et al. 2016; Proenca et al. 2017). *Land-Use* and *Land-Cover Change* (LUCC) due to agricultural expansion, hydropower development and pastureland intensification are among the most visible footprints of anthropogenic influence on the landscape (Rockstrom et al. 2009; Foley et al. 2011). A growing human population, increasing consumption and global food-security concerns have led to surges in production of agricultural and animal products (Godfray et al. 2010; Lambin and Meyfroidt 2011; Dias et al. 2016) that cause acute LUCC transformations in the tropics (Hansen et al. 2010).

Amazonian forests are home to nearly a quarter of the world’s known terrestrial biodiversity and are drivers of global atmospheric circulation (Malhi et al. 2008). As such, they are vital not only for biodiversity conservation but also for climate change mitigation (Pereira and Viola 2019). Nevertheless, humans continue to modify this biome since pre-Columbian occupation (Lombardo et al. 2013; Levis et al. 2017; McMichael et al. 2017). Although Amazonian deforestation rates appeared to be slowing in the first decade of the 21^st^ century (Nepstad et al. 2014), since then, they have continually increased to unprecedented levels (Escobar 2019a, Ferrante and Fearnside 2019) as Brazil’s recent policies have progressively weakened environmental laws in favor of the energy production and agribusiness sectors (Rajão and Georgiadou 2014; Brancalion et al. 2016; Fearnside 2016a; Abessa et al. 2019; Ferrante and Fearnside 2019; Scarrow 2019). Across the Amazon, the construction of roads that enable transport of timber (legal and illegal), deforestation for cattle ranching and the establishment of large monocultures (e.g., soy, corn, cotton) are the leading causes of Amazonian deforestation and deterioration of soil and water quality (Abell et al. 2011; Durigan et al. 2013; Macedo et al. 2013; Dias et al. 2015; Lessa et al. 2015; Lees et al. 2016; Marmontel et al. 2018; de Mello et al. 2018a; de Mello et al. 2018b; Rodrigues et al. 2018; Zimbres et al. 2018; Gauthier et al. 2019; Ilha et al. 2019; Klarenberg et al. 2019). In soybeans alone, Brazil increased its gross production by 94% between 1991 and 2013 (FAO 2016). The Brazilian Ministry of Agriculture estimated that grain and beef production will increase by nearly 30% by 2020 (MAPA 2015). Policies favoring agribusinesses have also deregulated the use of agrochemical products, many of which have been banned in other countries for years; 30% of agrochemical products used in Brazil are prohibited by the European Union (Costa et al. 2018). The widespread use of agrochemicals has considerably increased the contamination indexes of soil and water bodies across the country. These agrochemicals have been further shown to have deleterious effects on human health as well as biodiversity (Lugowska 2018; Martyniuk et al. 2020; Neves et al. 2020; Ribeiro et al. 2020; Velmurugan et al. 2020).

The Xingu River is the fourth largest tributary of the Amazon River, traversing ∼2,700 km through the Brazilian states of Mato Grosso and Pará. Its basin covers 510,000 km^2^ (51M ha), of which 200,000 km^2^ (20M ha) are Indigenous land and 80,000 km^2^ (8M ha) are under various levels of state or federal protection (Villas-Boas 2012). The Xingu watershed encompasses a complex geomorphological base (Sawakuchi et al. 2015; Silva et al. 2015) which promotes an extraordinary aquatic biodiversity and high rates of endemism (Sabaj 2015). Based on information compiled from published manuscripts and ichthyological collections, Dagosta and de Pinna (2019) found 502 valid species in the Xingu basin, with 13 of 367 species endemic to lower Xingu and 15 of 333 species endemic to the upper portion of the river. Taking the undescribed species into account, there are an estimated 600 species of fish, 10% of them endemic to the river basin (Camargo et al. 2004; Barbosa et al. 2015; Giarrizzo et al. 2015; Fitzgerald et al. 2018; Jézéquel et al. 2020a).

Cattle have been present on the Xingu floodplains for over 100 years, but prior to the 1970s the herds were relatively small, and deforestation was minimal (Goulding et al. 2003). Large herds of cattle were introduced in the 1980s. As a result, the natural forest outside of protected areas and Indigenous territories has been heavily deforested or modified (Goulding et al. 2003; Dias et al. 2016). The displacement of the cattle ranches by large-scale agriculture subsequently led to the greatest rates of deforestation in the region (Barona et al. 2010). Proliferation of genetically modified soy has further increased crop yields, and the Xingu basin provides 8% of the World’s soy production. Mega-scale soy plantations are considered as one of the main threats to the environment and rural livelihoods in Brazil (Fearnside 2001).

The Xingu basin also has an intricate cultural mosaic of Indigenous tribes and “ribeirinhos” living on the floodplain; continuous human occupation has modified the landscape over at least the last 1500 years (Schwartzman et al. 2013). In addition, over 200 ribeirinho settlements of various sizes from single family occupations to small communities are estimated to reside on the Xingu floodplains each with various sized footprints of deforestation and forest degradation (Schwartzman et al. 2013). Selective logging, the spatially diffuse thinning of large commercially valuable trees as opposed to clear-cutting, has also been shown to occur frequently throughout the Brazilian Amazon, and in the Xingu basin, both within and outside of protected areas and Indigenous lands (Asner et al. 2005).

While conservation priorities often neglect ichthyofauna (Stiassny 1996; Nogueira et al. 2010; Abell et al. 2011), some initiatives have begun to include freshwater ecosystems, recognizing the separate nature of aquatic and terrestrial diversity (Olson and Dinerstein 2002; Abell et al. 2008; Beaumont et al. 2011; Dinerstein et al. 2019). Quantifying change in a biologically and anthropogenically diverse landscape such as the Xingu basin is paramount. Given its large extent, land-cover products derived from remote sensing are the most efficient sources of such information. Importantly, these data also provide historical assessments of change for the last four decades. Several land cover products have been generated at varying scales and time periods by different agencies/groups. Despite individually reporting high levels of accuracy, uncertainty and disagreement between them has been documented (Foody 2004; Foody 2006; Herold et al. 2006; Fritz and See 2008), including limited agreement on the spatial distribution of the individual classes (McCallum et al. 2006; Tsendbazar et al. 2016). Therefore, despite acknowledgement of deforestation as a primary driver of biodiversity loss, uncertainty in baseline estimates persists (Sexton et al. 2016).

Here, our objective is to compare forest cover and surface water estimates over time for the Xingu basin from pre-existing remotely sensed classifications in order to determine potential impacts of land cover map choice on conservation of fish fauna. These datasets continue to be commonly used in ecological studies to for example, assess the impacts of Riparian forest loss on freshwater biodiversity (Dala-Corte et al. 2020), assess greenhouse gas fluxes from hydropower dam reservoirs (Araújo et al. 2019), define priority areas for ichthylogical conservation (Jézéquel et al. 2020b), define areas for forest restoration (Lopes et al. 2020), study the impacts of mining (Siqueira-Gay et al. 2020), and link illegal deforestation rural properties to agricultural production and exports (Rajão et al. 2020), among others, often without a justification for their use or critical assessment of their suitability or limitations.

There is a broad body of literature examining the importance of terrestrial land cover, and in particular, forest cover and its conversion, on the health and biodiversity of freshwater ecosystems (e.g. Gregory et al. 1991; Harding et al. 1998; Leal et al. 2017; Guimarães Frederico et al. 2018; Luke et al. 2018; Lamboj et al. 2020, among others). A key finding from a systematic review by Lo et al. (2020) on the importance of forests on freshwater fish communities was the consistent evidence that forests positively influence tropical fish diversity. Another aspect highlighted was the importance of forested areas for fish species with narrow habitat preferences as opposed to generalists that are abundant in disturbed areas. Changes in forest cover and fragmentation have a negative impact on aquatic species, many of them endemic to the Xingu basin. Furthermore, given the current politicization of conservation in Brazil (Abessa et al. 2019; Escobar 2019b; Fuchs et al. 2019), understanding differences between the many remotely sensed baselines is fundamentally important to avoid information misuse and to objectively indicate potential issues when used for conservation or policy making.

## MATERIAL AND METHODS

### Study area

The study area is ∼1.3 million km^2^ (130M ha) encompassing the Xingu river basin, Brazil (Figure 1). This area includes portions of five states (Amazonas, Pará, Mato Grosso, Tocantins and Goiás) and two diverse biomes (Amazonia and Cerrado). The Upper Xingu headwaters are spread across the sedimentary Parecis basin (Sabaj 2015). Many of these headwaters have little or no protection and are increasingly affected by industrial agriculture, leading to massive erosion of the riverbanks as well as contamination from pesticides and fertilizers (Sanches et al. 2012). Recent human activity has increased industrial agriculture (i.e. soy, cotton and corn) and extensive cattle ranches leading to deforestation and associated decrease in stream water quality (Barona et al. 2010; Durigan et al. 2013; de Mello et al. 2018b; Stella and Bendix 2019). A more reliable road system has also opened not only the agrarian frontier (Schmink et al. 2019) but has also facilitated artisanal illicit gold mining operations leading to elevated mercury contamination (Da Silva et al. 2018a). These headwaters are home to a number of endemic fishes (e.g., the cichlid *Apistogramma kullanderi*, the catfish *Harttia rondoni* and the characins *Moenkhausia petymbuaba, M. chlorophthalma* and *Knodus nuptialis*) with distributions limited to one or a few streams.

**Figure 1.**
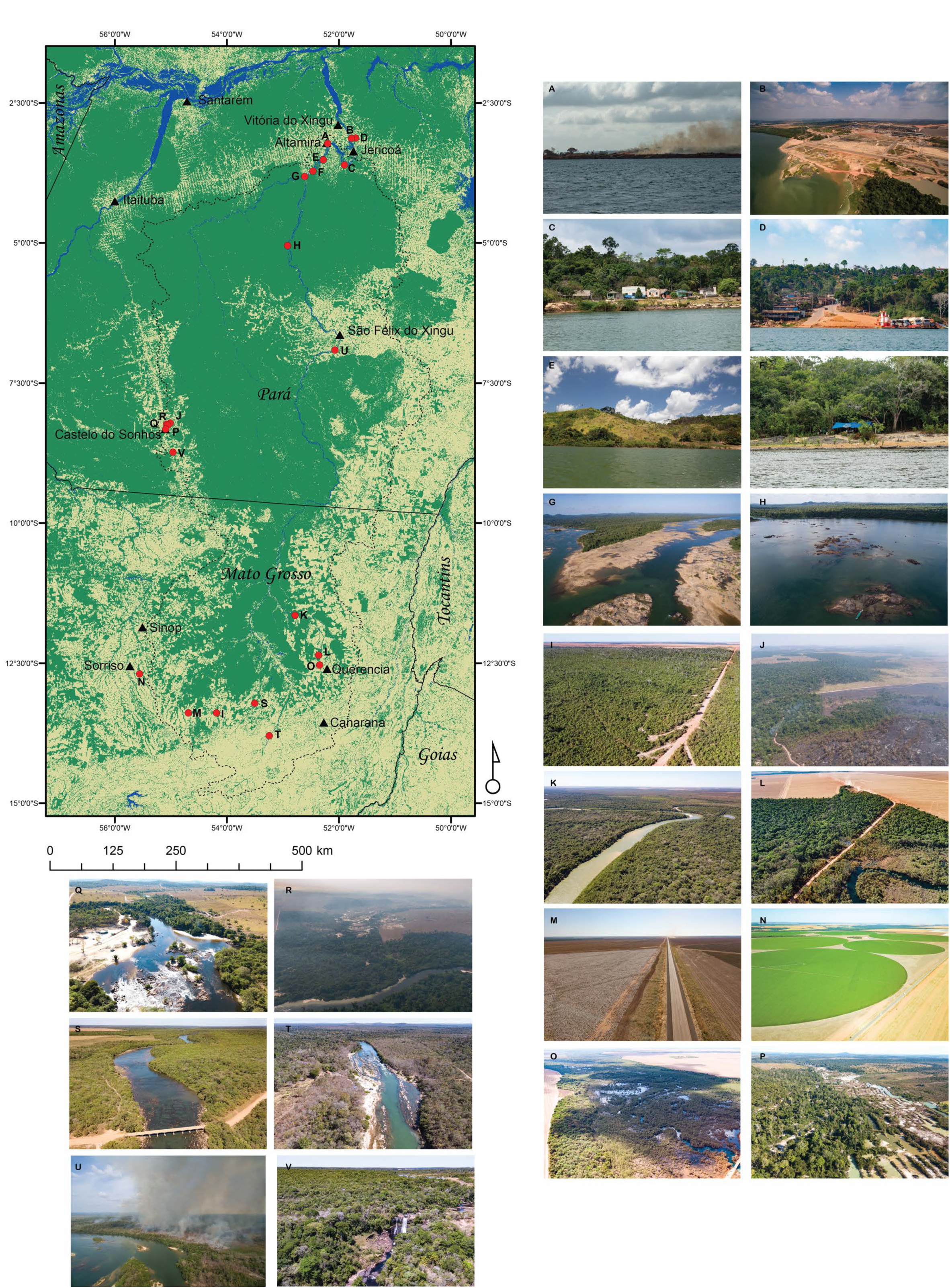
Map of study area with examples of natural and altered land cover. Background map is the Fish+Forest 2018 classification (Kalacska et al. 2019a) Dashed line illustrates the outline of the Xingu freshwater ecoregion from Abell et al. (2008). A) clearing and burning of Arapujá island across from Altamira, B) Belo Monte dam, C) gold mining settlement, D) urban area at the ferry crossing between Belo Monte I and Belo Monte II, E) forest clearing for pasture, F) small homestead, G) confluence of the Xingu and Iriri rivers in the dry season with substrate exposed, H) intact forest in protected area, I) new road construction, J) large scale burning, K) intact forest surrounded by agricultural fields, L) agricultural encroachment on wetlands, M) mega-scale cotton plantation, N) irrigated agriculture, O) Riparian forest and wetland surrounded by agriculture, P) illegal mining, Q) urban area, R) forest burning and influx of water contaminated by mine tailings, S) Riparian forest, T) deciduous forest surrounding Culuene rapids, U) large scale forest burning, V) intact forest surrounding Curuá rapids with the Hydro Energia Buriti dam reservoir in the background. An interactive 3D reconstruction of the landscape shown in T (http://bit.ly/culuene) is available from (Kalacska et al. 2019b). The interactive 3D reconstruction of V can be found at (http://bit.ly/altocurua).

The middle portion of the Xingu River drains Pre-Cambrian rocks of the Brazilian Shield and extends from São Félix do Xingu downstream to Belo Monte. The lowermost stretch of the middle Xingu, approximately 173.7 km (navigable river) from Altamira to Belo Monte, is known as the Volta Grande (Big Bend). The ichthyofauna of the middle Xingu differs greatly from that of its headwaters. The high levels of fish diversity and endemism (e.g. the loricariid catfish *Hypancistrus zebra, Baryancistrus xanthellus, Pseudacanthicus pirarara*, the serrasalmid *Ossubtus xinguense* and the cichlids *Crenicichla dandara, Teleocichla preta*, and *Cichla melaniae)* are due to its complex geomorphology. A 193.9 km distance (dry season navigable river) has been impacted by the Belo Monte Project, the second largest dam complex in Brazil and fourth largest in the world by installed capacity. Belo Monte operates on a run-of-river basis with electricity generation varying according to river flow. It continues a global trend of damming tropical rivers (Fearnside 2016a; Lees et al. 2016; Winemiller et al. 2016; Latrubesse et al. 2017). Major alterations to the river include the inundation of a minimum of 426 km^2^ (during low water) over a 62.7 km distance (Kalacska et al. 2020) of the main channel above the Pimental diversion dam, and dewatering of 131.2 km (low water navigable channel distance) between Pimental and the outlet for the Belo Monte dam where water is returned to the Xingu River. Highly specialized fishes and other aquatic organisms adapted for life in rheophilic habitats (e.g., Lujan & Conway 2015) are not able to survive under the new conditions. Also, almost all aquatic species rely on the river’s annual flood pulse to complete their life cycle (Junk et al. 1989). The Xingu’s natural flood pulse averaged 4.8 m per year (Barbosa et al. 2015). Artificial regulation of the Xingu’s water levels fails to mimic the natural flood pulse (Figure 2) and has already had devastating effects on the ichthyofauna of Volta Grande (Souza-Cruz-Buenaga et al. 2019). In addition, in the section of the Volta Grande dewatered by the Belo Monte Dam, there is yet another project, called “Projeto Volta Grande” planned by the Belo Sun mining company, to begin massive gold extraction in an already weakened ecosystem (Tófoli et al. 2017; Bratman and Dias 2018).

**Figure 2.**
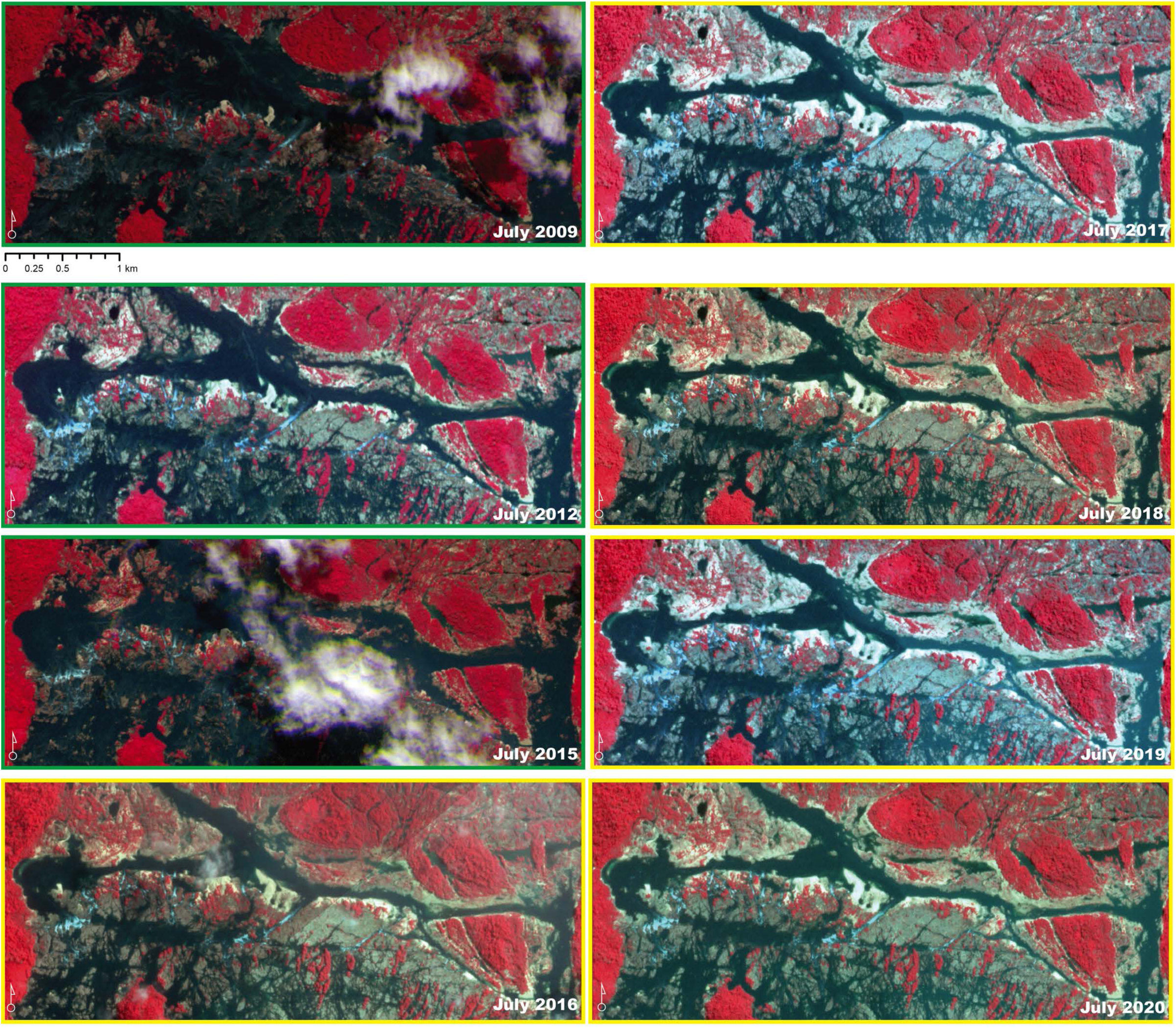
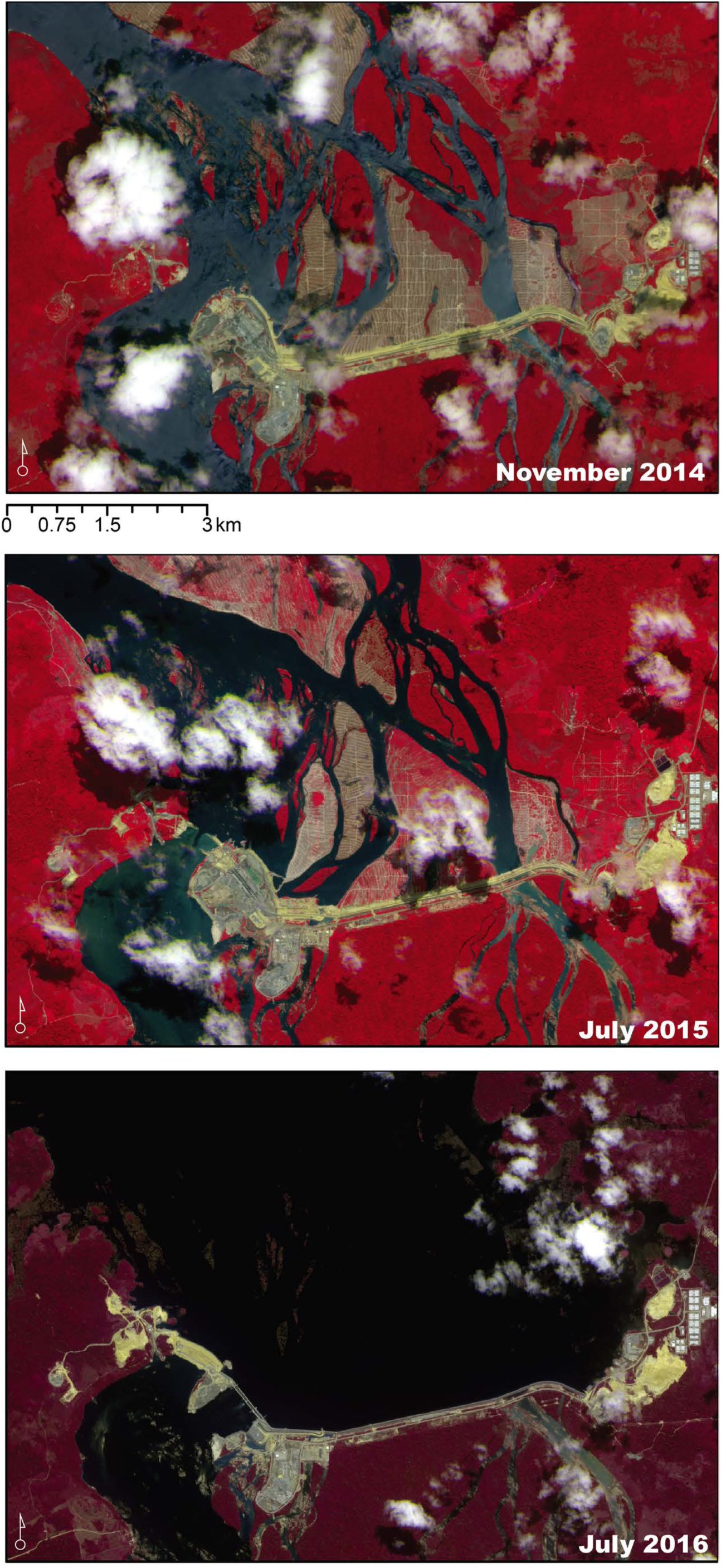
A) Comparison of the Xingu river water level at the Jericóa rapids before (green border) and after (yellow border) operationalization of Belo Monte. Images are from the RapidEye constellation (5 m pixels) for 2009 - 2015 and the PlanetScope constellation (3 m pixels) for 2016 - 2020. All images are shown as near infrared/red/green composites to more clearly differentiate water from the exposed rocks. A consistently lower water level can be seen in all images post operationalization. B) Imagery from the PlanetScope constellation of the construction of the Pimental dam. All images are shown in as a near infrared/red/green composite to more clearly differentiate water and the progress of the construction.

The lower portion of the Xingu where it enters the Amazonas sedimentary basin is referred to as the Xingu Ria. The Xingu Ria is home to different fish fauna comprised of both cosmopolitan species found throughout the Amazon lowlands (e.g., the piraiba *Brachyplatystoma filamentosum*) including a few species that move upstream into the Volta Grande (e.g., the redtail catfish *Phractocephalus hemioliopterus*), and endemic species such as *Hypancistrus* sp. “acari pão” and the recently described *Panaqolus tankei*.

### Remotely sensed datasets

We downloaded eleven commonly used, freely available regional and global land cover and surface water classifications created from remotely sensed data (Tables 1 and 2). We chose four time periods for which multiple datasets were available: 1989-1992, 2000, 2010, 2014-2018. Because each dataset was created from different imagery with varying methodologies, and have different output characteristics (e.g., spatial resolution, definition of forest) we evaluated each dataset in context to illustrate their similarities and differences. The spatial resolutions vary from 25 m (JAXA Global Forest / Non-forest; Shimada et al. 2014) to 300 m (ESA Land Cover CCI; Defourny et al. 2016) and include both optical and radar satellite sensors (Tables 1 and 2). A comprehensive review of sensor characteristics and remote sensing concepts for aquatic applications can be found in Rowan and Kalacska (2020), however, Tables 3, 4 and Figure 3 provide a description and illustration of the main remote sensing concepts addressed in this study.

**Table 1.** Summary of land cover products used in the comparisons. ^*^Tree cover threshold of greater or equal to 30% applied. ^**^The observational unit is 81 ha (Defourny et al. 2016). Data products are created from “passive sensors” relying on reflected solar radiation with the exception of the two marked with a ^§^ which are from active radar. ^†^15 sec resolution at the equator is 463.8 m longitude x 460.7 m latitude. At 10°S it becomes 456.8 m longitude x 460.9 m latitude.

| Map | Year(s) | Sensor(s) | Pixel size (m) | MMU (ha) | Classes | Reference |
| --- | --- | --- | --- | --- | --- | --- |
| ESA | 1992, 2000, 2010, 2015 | MERIS FR/RR, AVHRR, SPOT-VG, PROBA-V | 300 | 9** | 22 land cover classes | Bontemps et al. (2013) ; Kirches et al. (2014) |
| EC JRC/Google | 1984-2015 and 1984-2018 | Landsat TM5, 7 ETM+, OLI 8 | 30 | N/A | Surface water occurrence, maximum extent | Pekel et al. (2016) |
| Fish + Forest | 1989, 2000, 2010, 2018 | Landsat TM5, OLI 8 | 30 | 0.8 | Forest, Non-forest, Water | Kalacska et al. (2019a) |
| Global Forest Change* | 2000 | Landsat 7 ETM+ | 30 | N/A | percent tree cover | Hansen et al. (2013) |
| JAXA | 2010, 2017 | PALSAR§/PALSAR-2§ | 25 | 0.5 | Forest, Non-forest, Water | Shimada et al. (2014) |
| HydroSHEDS | 2000 | Shuttle Radar Topography Mission (SRTM) | 15 sec† | N/A | Water | Lehner et al. (2008) |
| MapBiomass | 1989, 2000, 2010, 2017 | Landsat TM5, 7 ETM+, OLI 8 | 30 | 0.5 | 27 Land cover classes | http://mapbiomas.org |
| MODIS Vegetation Continuous Fields * | 2000, 2010, 2015 | MODIS | 250 | N/A | percent tree cover | Dimiceli et al. (2015) |
| PRODES | 2010, 2017 | Landsat, MODIS, Terra, CBERS, ResourceSat | 90, 30 | 6.25 | 5 land cover classes and 19 years of deforestation | INPE (2015); Rajão et al. (2017) |
| TanDEM-X Global Forest/Non-Forest | 2016 | TanDEM-X<sup>§</sup> | 50 | 0.5 | Forest, non-forest, water | Esch et al. (2017); Martone et al. (2018) |
| Terra Class Amazônia and Cerrado | 2010, 2014 | Landsat, MODIS, SPOT-5 | 30 | 6.25 | 15-16 land cover classes | de Almeida et al. (2016); Rajão et al. (2017) |

**Table 2.**
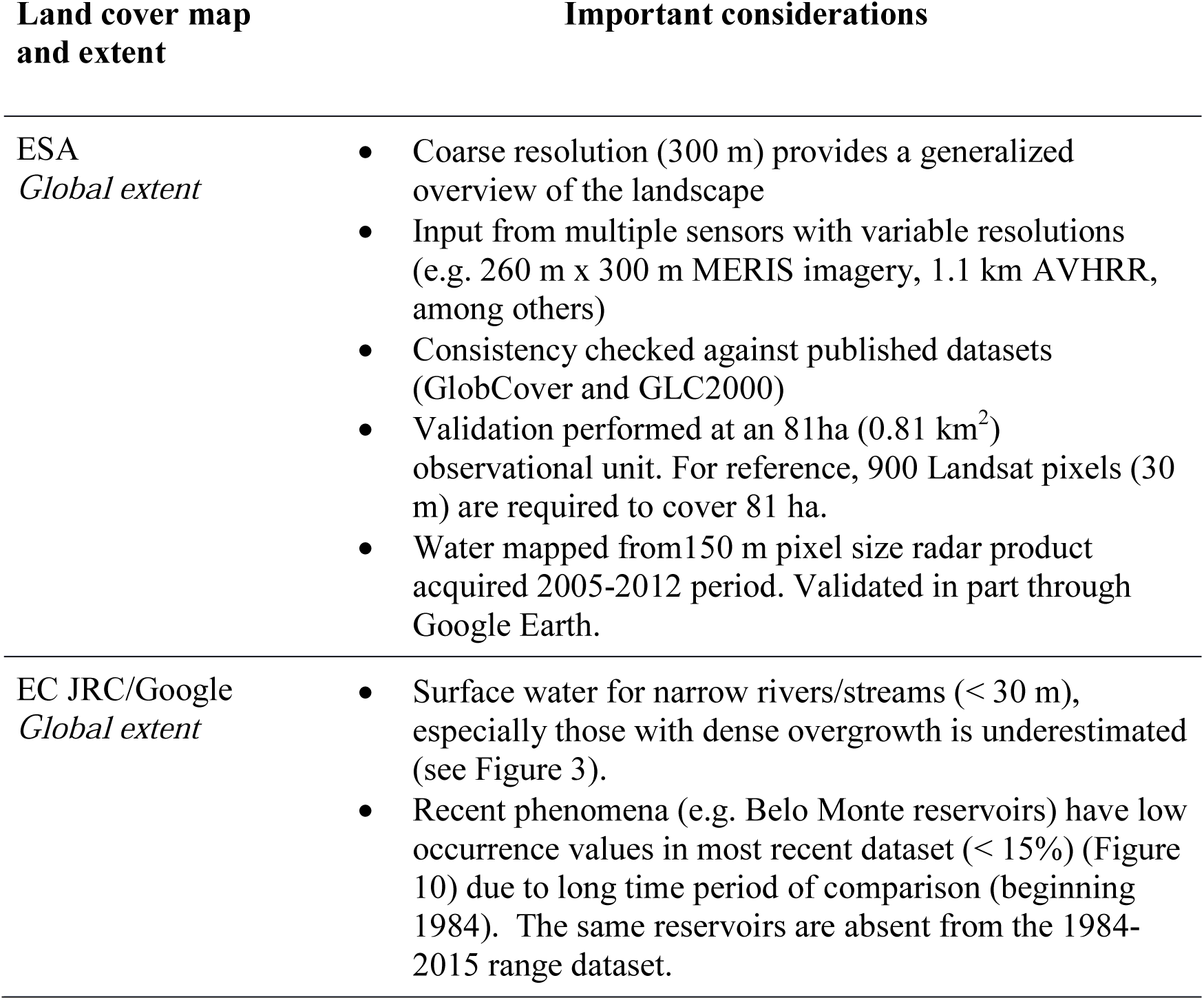
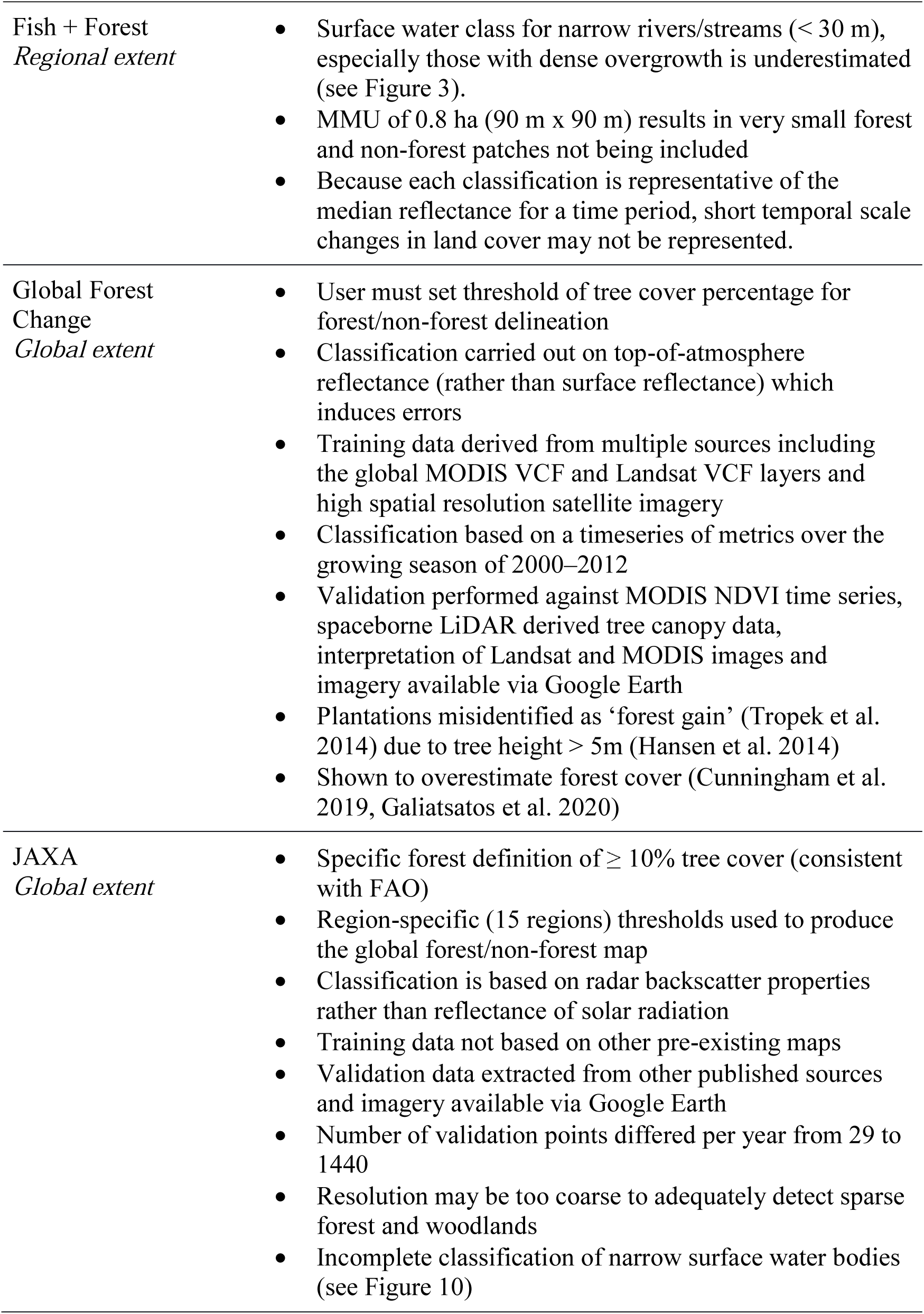
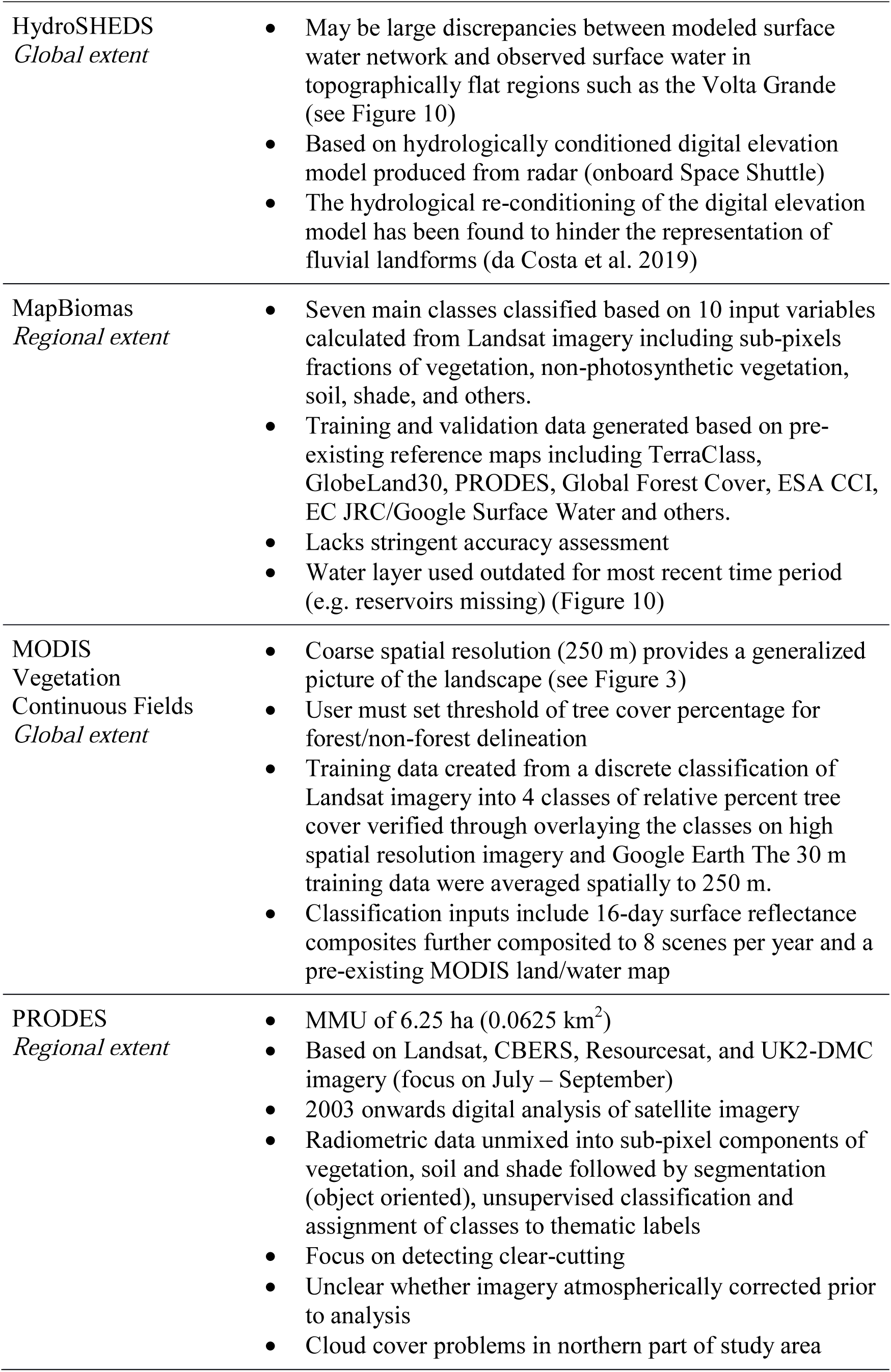
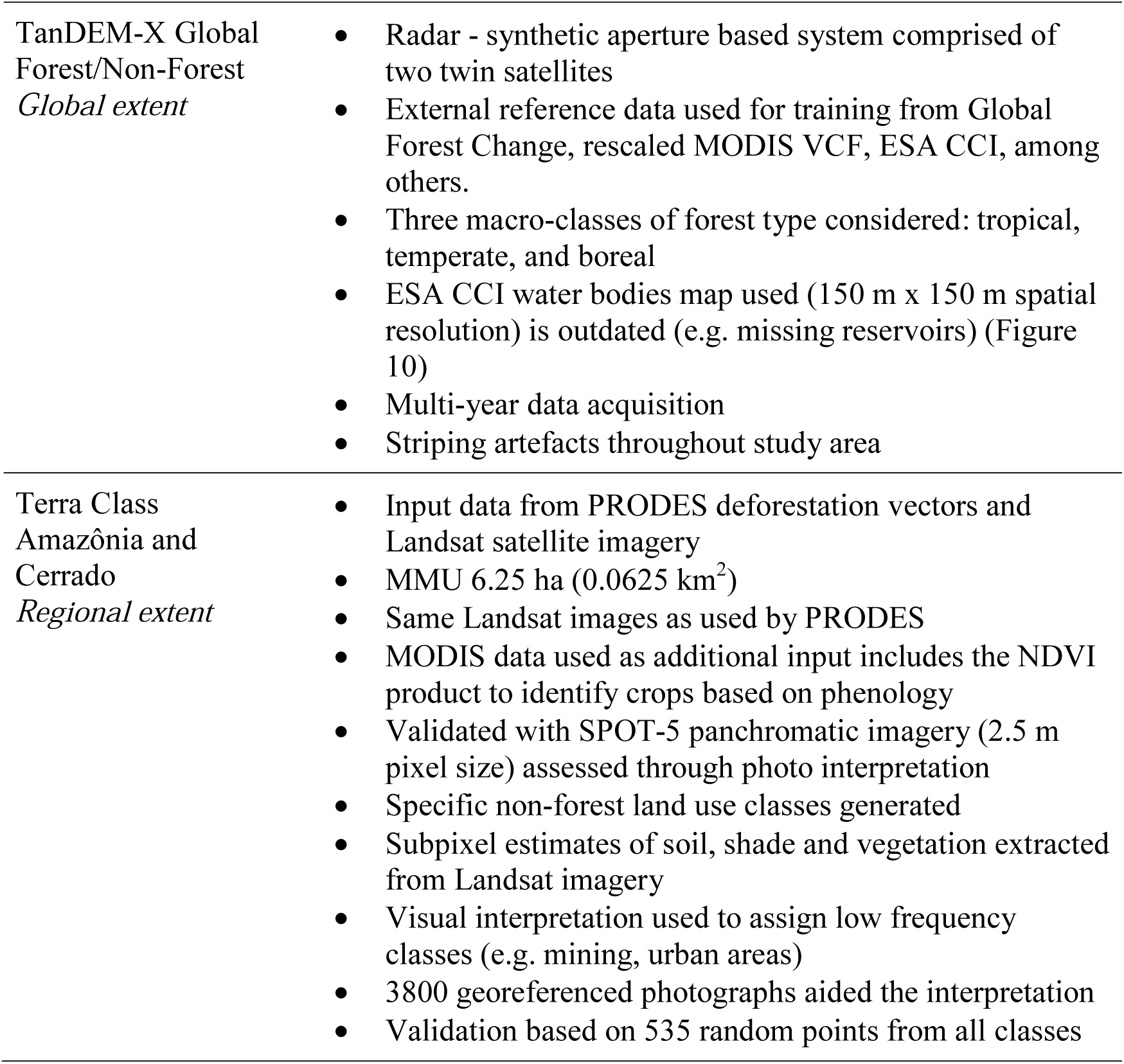
Summary of important considerations and limitations of the land cover products used in the comparisons.

| Land cover map and extent | Important considerations |
| --- | --- |
| ESA<br>*Global extent* | • Coarse resolution (300 m) provides a generalized overview of the landscape<br>• Input from multiple sensors with variable resolutions (e.g. 260 m x 300 m MERIS imagery, 1.1 km AVHRR, among others)<br>• Consistency checked against published datasets (GlobCover and GLC2000)<br>• Validation performed at an 81ha (0.81 km<sup>2</sup>) observational unit. For reference, 900 Landsat pixels (30 m) are required to cover 81 ha.<br>• Water mapped from 150 m pixel size radar product acquired 2005-2012 period. Validated in part through Google Earth. |
| EC JRC/Google<br>*Global extent* | • Surface water for narrow rivers/streams (< 30 m), especially those with dense overgrowth is underestimated (see Figure 3).<br>• Recent phenomena (e.g. Belo Monte reservoirs) have low occurrence values in most recent dataset (< 15%) (Figure 10) due to long time period of comparison (beginning 1984). The same reservoirs are absent from the 1984-2015 range dataset. |
| Fish + Forest<br>*Regional extent* | • Surface water class for narrow rivers/streams (< 30 m), especially those with dense overgrowth is underestimated (see Figure 3).<br>• MMU of 0.8 ha (90 m x 90 m) results in very small forest and non-forest patches not being included<br>• Because each classification is representative of the median reflectance for a time period, short temporal scale changes in land cover may not be represented. |
| Global Forest Change<br>*Global extent* | • User must set threshold of tree cover percentage for forest/non-forest delineation<br>• Classification carried out on top-of-atmosphere reflectance (rather than surface reflectance) which induces errors<br>• Training data derived from multiple sources including the global MODIS VCF and Landsat VCF layers and high spatial resolution satellite imagery<br>• Classification based on a timeseries of metrics over the growing season of 2000–2012<br>• Validation performed against MODIS NDVI time series, spaceborne LiDAR derived tree canopy data, interpretation of Landsat and MODIS images and imagery available via Google Earth<br>• Plantations misidentified as ‘forest gain’ (Tropek et al. 2014) due to tree height > 5m (Hansen et al. 2014)<br>• Shown to overestimate forest cover (Cunningham et al. 2019, Galiatsatos et al. 2020) |
| JAXA<br>*Global extent* | • Specific forest definition of $\geq 10\%$ tree cover (consistent with FAO)<br>• Region-specific (15 regions) thresholds used to produce the global forest/non-forest map<br>• Classification is based on radar backscatter properties rather than reflectance of solar radiation<br>• Training data not based on other pre-existing maps<br>• Validation data extracted from other published sources and imagery available via Google Earth<br>• Number of validation points differed per year from 29 to 1440<br>• Resolution may be too coarse to adequately detect sparse forest and woodlands<br>• Incomplete classification of narrow surface water bodies (see Figure 10) |
| HydroSHEDS<br>*Global extent* | • May be large discrepancies between modeled surface water network and observed surface water in topographically flat regions such as the Volta Grande (see Figure 10)<br>• Based on hydrologically conditioned digital elevation model produced from radar (onboard Space Shuttle)<br>• The hydrological re-conditioning of the digital elevation model has been found to hinder the representation of fluvial landforms (da Costa et al. 2019) |
| MapBiomass<br>*Regional extent* | • Seven main classes classified based on 10 input variables calculated from Landsat imagery including sub-pixels fractions of vegetation, non-photosynthetic vegetation, soil, shade, and others.<br>• Training and validation data generated based on pre-existing reference maps including TerraClass, GlobeLand30, PRODES, Global Forest Cover, ESA CCI, EC JRC/Google Surface Water and others.<br>• Lacks stringent accuracy assessment<br>• Water layer used outdated for most recent time period (e.g. reservoirs missing) (Figure 10) |
| MODIS<br>Vegetation<br>Continuous Fields<br>*Global extent* | • Coarse spatial resolution (250 m) provides a generalized picture of the landscape (see Figure 3)<br>• User must set threshold of tree cover percentage for forest/non-forest delineation<br>• Training data created from a discrete classification of Landsat imagery into 4 classes of relative percent tree cover verified through overlaying the classes on high spatial resolution imagery and Google Earth The 30 m training data were averaged spatially to 250 m.<br>• Classification inputs include 16-day surface reflectance composites further composited to 8 scenes per year and a pre-existing MODIS land/water map |
| PRODES<br>*Regional extent* | • MMU of 6.25 ha (0.0625 km<sup>2</sup>)<br>• Based on Landsat, CBERS, Resourcesat, and UK2-DMC imagery (focus on July – September)<br>• 2003 onwards digital analysis of satellite imagery<br>• Radiometric data unmixed into sub-pixel components of vegetation, soil and shade followed by segmentation (object oriented), unsupervised classification and assignment of classes to thematic labels<br>• Focus on detecting clear-cutting<br>• Unclear whether imagery atmospherically corrected prior to analysis<br>• Cloud cover problems in northern part of study area |
| TanDEM-X Global Forest/Non-Forest<br>*Global extent* | • Radar - synthetic aperture based system comprised of two twin satellites<br>• External reference data used for training from Global Forest Change, rescaled MODIS VCF, ESA CCI, among others.<br>• Three macro-classes of forest type considered: tropical, temperate, and boreal<br>• ESA CCI water bodies map used (150 m x 150 m spatial resolution) is outdated (e.g. missing reservoirs) (Figure 10)<br>• Multi-year data acquisition<br>• Striping artefacts throughout study area |
| Terra Class Amazônia and Cerrado<br>*Regional extent* | • Input data from PRODES deforestation vectors and Landsat satellite imagery<br>• MMU 6.25 ha (0.0625 km<sup>2</sup>)<br>• Same Landsat images as used by PRODES<br>• MODIS data used as additional input includes the NDVI product to identify crops based on phenology<br>• Validated with SPOT-5 panchromatic imagery (2.5 m pixel size) assessed through photo interpretation<br>• Specific non-forest land use classes generated<br>• Subpixel estimates of soil, shade and vegetation extracted from Landsat imagery<br>• Visual interpretation used to assign low frequency classes (e.g. mining, urban areas)<br>• 3800 georeferenced photographs aided the interpretation<br>• Validation based on 535 random points from all classes |

**Table 3.** Definitions of main remote sensing concepts, from an Earth observation perspective, mentioned in this study. Readers are referred to fundamental reference texts such as Schowengerdt (2007), Lillesand et al. (2008), Jones and Vaughn (2010) and Manolakis et al. (2016) for further detail.

| Concept | Definition |
| --- | --- |
| Classification | Grouping pixels based on common attributes. |
| Ground Instantaneous Field of View (GIFOV) | Projection of the IFOV onto the ground surface. The GIFOV is generally larger than the pixel size/resolution. (See Figure 3) |
| Instantaneous Field of View (IFOV) | Angle over which an individual element on a sensor is responsive. (See Figure 3) |
| LiDAR | Acronym for *Light Detection And Ranging*. Active sensor system that uses a pulsed laser to measure distances to objects and/or the surface of the Earth. |
| Multispectral Imaging | Generally, refers to a digital image where each pixel is comprised of 3-15 spectral bands/layers. |
| Nadir | The point directly below the observer/observation point. Looking straight down. |
| Optical imagery | Images acquired by recording the energy reflected from materials within the optical portion of the electromagnetic spectrum which covers the visible to shortwave infrared (400-2500 nm) range. |
| Pixel size | Also referred to as ground sampling distance (GSD) in some sources. It is the distance between pixels (also equivalent to pixel width). It comprises the majority of the area on the ground contributing signal to a pixel (see Figure 3). Most often used to describe an image after it has been geometrically corrected and resampled (to square pixels) but can also refer to the raw data with unaltered geometry (see Inamdar et al. 2020 for an example). Importantly, neighboring pixels are not independent |
| Radar data | Acronym for *Radio Detection And Ranging*. Most commonly used radar systems for Earth observation are active sensors that generate their own signal to illuminate the target. These microwave sensors operate at longer wavelengths (~1 cm – 1m) than optical sensors. |
| Raster | A continuous data model that is a digital matrix of rows and columns. Each cell within the matrix is a pixel and contains a digital number related to the parameter acquired by the sensor. |
| Spatial resolution | The smallest resolvable detail in a photograph or image. Refers to the differentiation between two closely spaced objects, not the size of the actual objects. |
| Spectral resolution | Width of the wavelength interval for each band/layer of an image. |
| Temporal resolution | Revisit time of acquired data over the same area/scene (see Table 4). |
| Training data | A subset of example data representing classes of interest (e.g. forest, water, soil) used to develop (i.e. train) classification models. |
| Validation data | An independent subset of data used to assess the accuracy of a classification. |
| Vector | A discrete data model that uses points, lines and polygons to represent objects on the surface of the Earth. |

**Table 4.**
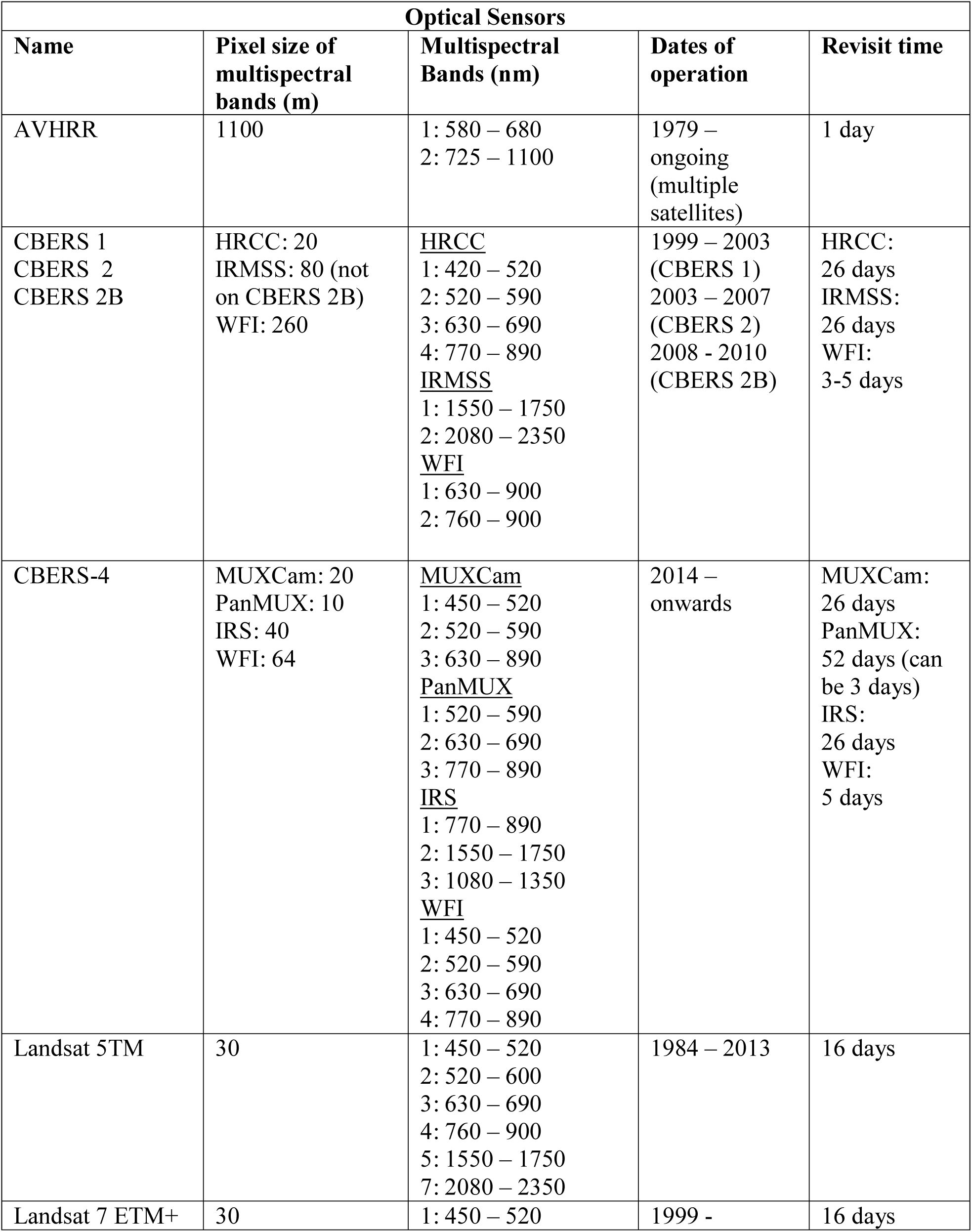
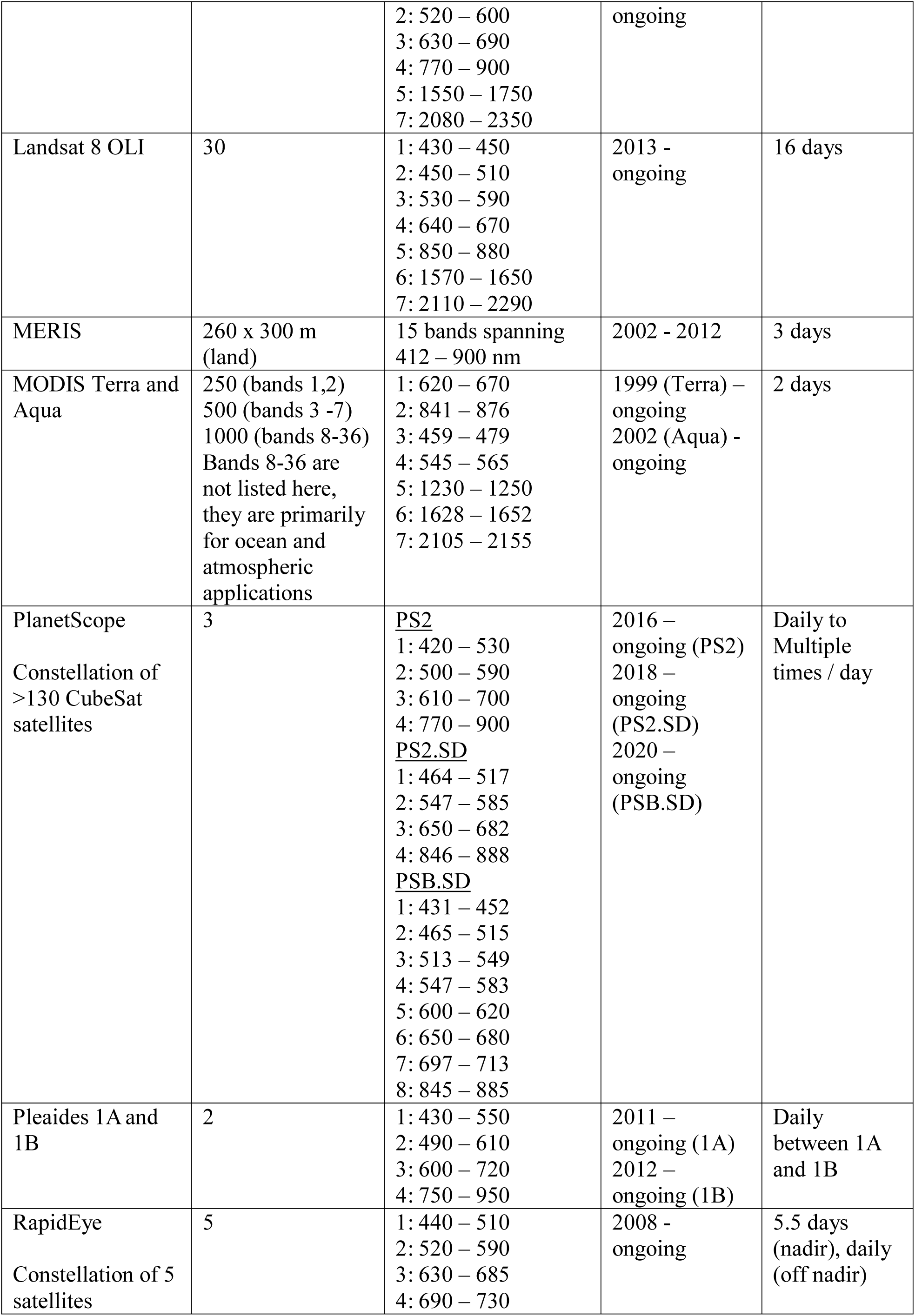
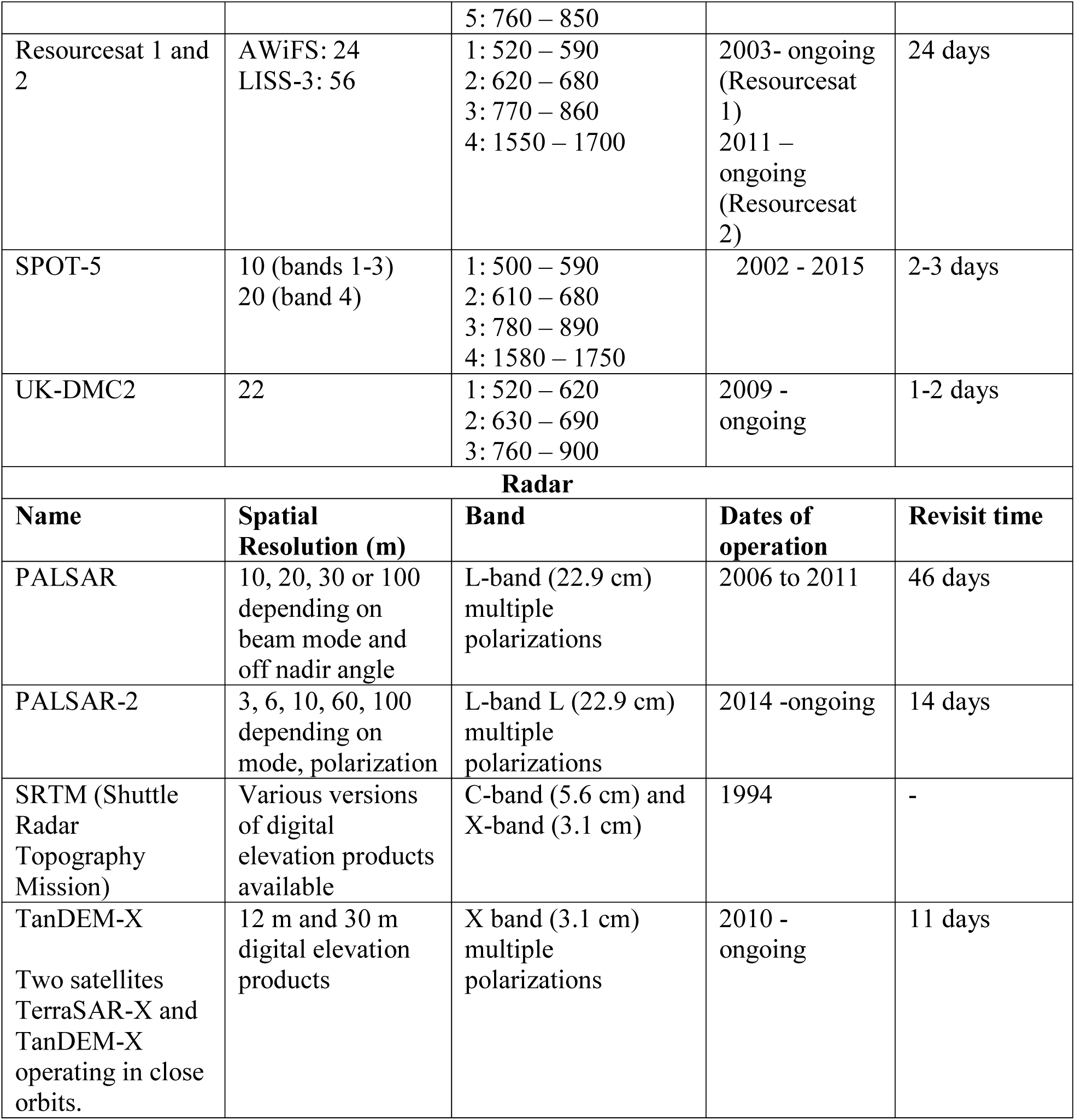
Main characteristics of spaceborne sensors referred to in this study. In addition to the multispectral channels described here, many of the optical sensors listed also have panchromatic, thermal or other specialized bands with different spatial and spectral characteristics.

**Figure 3.**
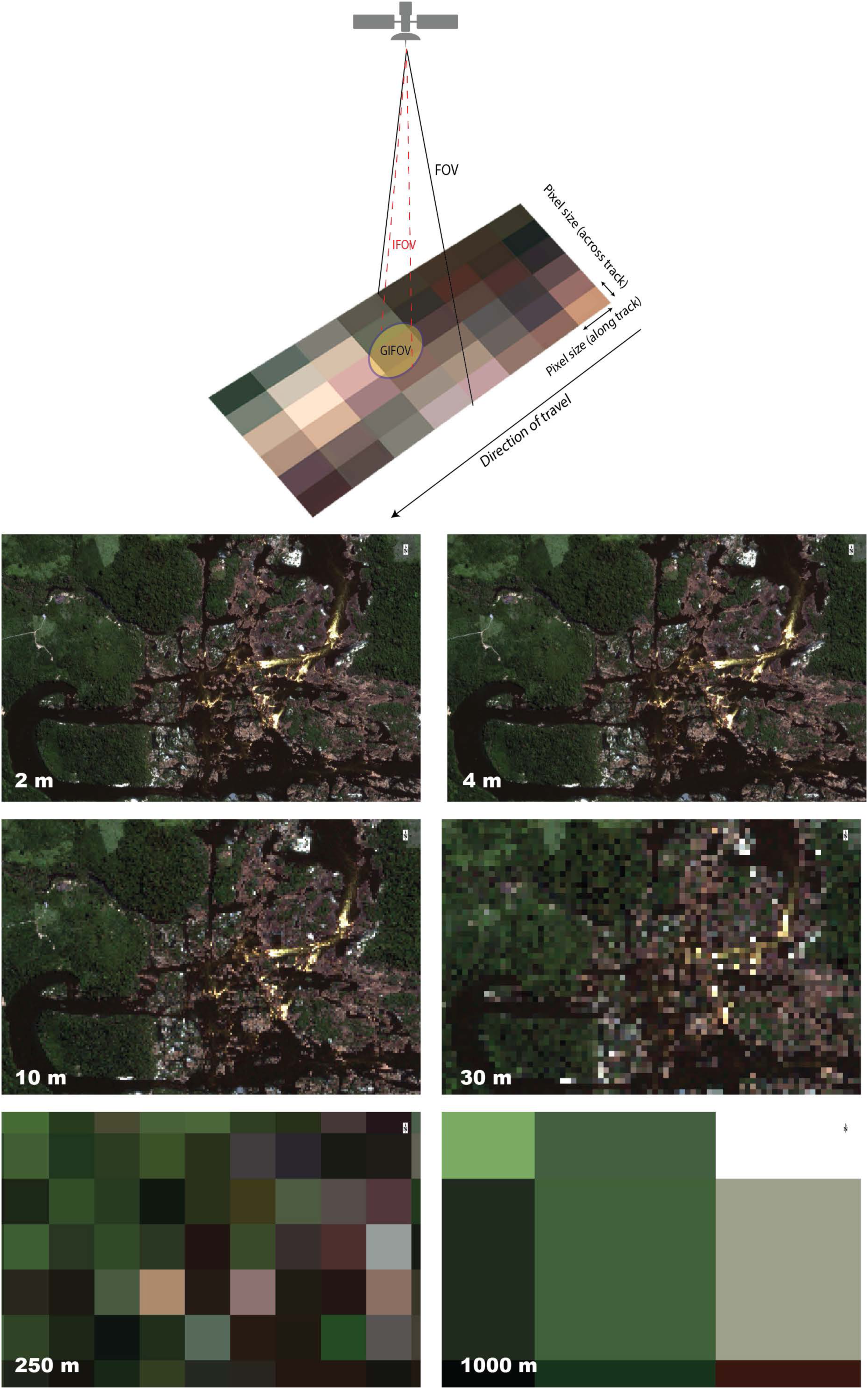
Diagram shows the important spatial concepts described in Table 3. Of note, the GIFOV spans more than one pixel leading to overlap between neighboring pixels. Also, pixels acquired by the majority of Earth observation optical sensors are not perfectly square in their raw geometry due to the motion of the sensor during acquisition. Pixels are most commonly represented as squares following geocorrection and resampling. See Inamdar et al. (2020) for additional details. Images subsets are an illustration of different pixel sizes for the differentiation of land covers suchs water, rocks and vegetation. The original Pleides 1A multispectral image acquired August 2, 2012 of Cachoeira Grande where the middle Xingu meets the Xingu Ria sectors was resampled from 2 m to 4 m, 10 m, 30 m, 250 m and 1000 m. Landsat imagery used in the majority of the classifications assessed in this study is provided at 30 m.

### Optical imagery

The Terra MODIS Vegetation Continuous Fields (VCF) (Dimiceli et al. 2015) and Global Forest Change (Hansen et al. 2013) classifications provide pixels values of percent tree cover at 250 m and 30 m spatial resolutions respectively. For both, we applied a threshold of ≥ 30% consistent with the Brazilian forest definition (Gilbert 2009). The other classification generated from optical imagery are provided with thematic pixels values representing forest and other classes (e.g. water). Generated yearly, the Terra MODIS VCF product is a sub-pixel-level estimation of surface vegetation cover. It is produced from monthly composites of Terra MODIS 250- and 500-meter resolution surface reflectance imagery and land surface temperature (Dimiceli et al. 2015). The Global Forest Cover baseline map (Hansen et al. 2013) was generated from cloud free mosaics of growing season Landsat 7 ETM+ images for the year 2000. Training data were generated from high spatial resolution Quickbird satellite imagery, as well as pre-existing vegetation continuous field maps derived from Landsat and MODIS imagery (Hansen et al. 2003; Hansen et al. 2011).

The ESA CCI Land Cover product (300 m spatial resolution) was generated for three reference periods each spanning five years to minimize cloud cover (Bontemps et al. 2013). The primary data sources for these classifications are MERIS Full Resolution (FR) images supplemented with MERIS Reduced Resolution (RR) and SPOT-VGT imagery. Consistency with reference datasets such as GlobCover (Arino et al. 2008) and GLC2000 (European Commission 2003) was verified in the preparation of the land cover maps. Validation was performed at an observational unit of 81 ha with samples derived from Google Earth, Landsat TM or ETM+ imagery and multi-temporal SPOT-VG indices.

The Fish + Forest classifications were created from cloud free mosaics of Landsat TM5 and Landsat 8 OLI imagery (Kalacska et al. 2019a). Due to extensive seasonal cloud cover in the region, each mosaic represented the median cloud free reflectance over a reference period of 2-5 years. Covering the smallest overall area of all the datasets examined, training data were generated based on field visits throughout the study area. No other pre-existing datasets were used in the classification process.

PRODES is a yearly product for the Amazon to monitor clear-cutting. It is one of eight forest monitoring systems developed by the Brazilian government (Rajão and Georgiadou 2014; INPE 2015; Rajão et al. 2017). TerraClass utilizes deforestation vector data from PRODES supplemented with satellite imagery from Landsat and MODIS (validated by SPOT-5 imagery) to monitor LUCC in the Brazilian Legal Amazon (de Almeida et al. 2016). MapBiomas classifications are yearly land cover products generated from Landsat imagery (MapBiomas Project). The products are developed separately by biome; here we include subsets for the Amazon and Cerrado. For the MapBiomas data, model training and validation samples are generated from reference maps including TerraClass, PRODES, Global Forest Cover, ESA CCI and others.

### Radar data

The TanDEM-X global forest classification (50 m spatial resolution) was generated from bistatic interferometric synthetic aperture radar (InSAR) data (HH polarization, X-band) acquired over the 2011-2015 period for the generation of a global digital elevation model (DEM) (Esch et al. 2017; Martone et al. 2018). Forest is defined as > 60% tree cover from the Global Forest Change map (Hansen et al. 2013). The JAXA global forest classification (25 m spatial resolution) is a yearly product (Shimada et al. 2014) generated from HH and HV polarized L-band Synthetic Aperture Radar (PALSAR and PALSAR-2). To be consistent with the FAO definition, areas with a tree cover ≥ 10% are included in the JAXA global forest dataset.

### Surface water

While a surface water class was included in eight of the landcover classifications described above, we also examined the EC JRC/Google (Pekel et al. 2016) product created exclusively for surface water (1984 – 2015 and 1984 – 2018 periods). We include the percent occurrence and maximum extent layers. We also include the HydroSHEDS river network. This is a model of the world’s rivers derived from the Shuttle Radar Topography Mission (SRTM) GL3 DEM. The network has been hydrologically conditioned using a sequence of automated procedures including sink filling, stream burning, and manual corrections, among others (Lehner et al. 2008). This dataset is different from the others because water occurrence is not directly observed, the river network is derived from the hydrologically conditioned DEM. It is one of the most commonly used base layers in ichthyological studies.

### Analyses

Each dataset was subset to the ∼1.3 million km^2^ area of interest (Figure 1) and total forest and water (where available) areas were calculated. A fragmentation analysis was carried out, examining four main class metrics for forest cover and their change over time for the different land cover classifications with FragStats v4 (McGarigal et al. 2012). ‘Class area’ (CA) and ‘percentage of landscape’ (PLAND) are important measures of landscape composition; specifically, how much of the landscape is comprised of a particular land cover (McGarigal et al. 2012). In this study, CA is a good indicator of the total forest area lost over time, while PLAND allows for comparison between maps of the percent of forest lost over time. For the three classifications available for all four time periods (i.e., ESA, Fish + Forest and MapBiomas), two additional indices were calculated: Largest Patch Index (LPI) and the Landscape Division Index (DIVISION). LPI measures the change in the landscape comprised by the largest patch, while DIVISION is a standardized measure of the probability of two randomly selected pixels in the landscape not being located in the same patch of the corresponding patch type (McGarigal et al. 2012). DIVISION ranges from 0 – 1, with a value of 0 when the landscape is comprised of a single patch. Due to a software limitation (32-bit software), the 25 m and 30 m spatial resolution datasets were resampled to 90 m prior to the calculation of these fragmentation metrics.

Spatial fragmentation analyses consisting of Entropy and Foreground Area Density (FAD) were calculated for each dataset with the GUIDOS Toolbox (Vogt and Riitters 2017). Spatial entropy is the degree of disorder in the spatial arrangement of the forest class. A single compact forest patch has minimum entropy whereas maximum entropy is found when the forest class is split into the maximum number of patches given the area and dispersed equally (i.e., checkerboard pattern; Vogt and Riitters 2017). FAD measures the proportion of forest pixels at five observation scales (7, 13, 27, 81, 243 pixels). Results are classified into six classes: < 10% (rare), 10-39% (patchy), 40-59% (transitional), 60-89% (dominant), 90-99% (interior), and 100% (intact). The final maps represent the average from the five observation scales (Vogt and Riitters 2017). Morphological Spatial Pattern Analysis (MSPA; Soille and Vogt 2009) was applied to each forest classification to map morphometric details of the forest patches. Of the seven mutually exclusive classes of core, islet, loop, bridge, perforation, edge, and branch mapped by MSPA for forest in each dataset we extracted the edge (external object perimeter) and core (interior excluding the perimeter) classes for further analysis.

## RESULTS

The forest cover maps for the four time periods are illustrated in Figure 4. As expected, the overall trend across datasets is a loss in forest area over time. Differences among the maps can be attributed to several factors including spatial resolution, cloud cover, input data, data quality, and methodology. For example, the forest northeast of Vitória do Xingu is noticeably absent in the PRODES 2010 map likely due to cloud cover. Also, the two coarse datasets (ESA and MODIS) have a more generalized representation of the forest cover. In addition, even within time periods there is a both an area and spatial arrangement difference in forest cover between datasets. For the earliest time period, the regions south and east of Canarana, as well as north of Santarém, have greater forest cover in the MapBiomas map than in either the ESA or Fish + Forest maps (Figure 4a). The same pattern can be seen for the other three time periods, as well with the MapBiomas data showing the greatest forest cover in that region. For the year 2000, the Global Forest Cover map is most similar to MapBiomas showing greater forest cover in the southern and eastern sections of the study area than the other classifications (Figure 4b).

**Figure 4.**
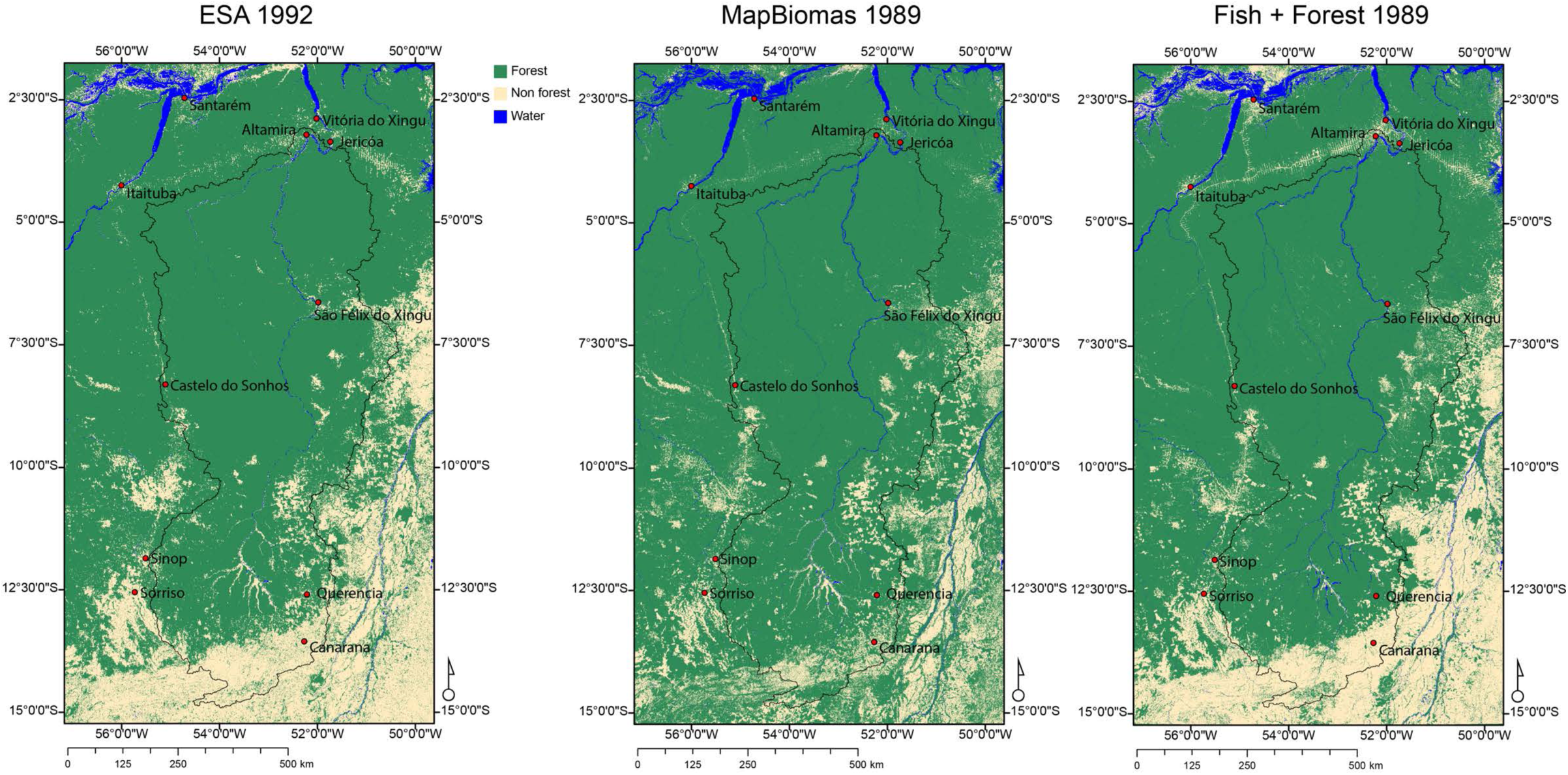
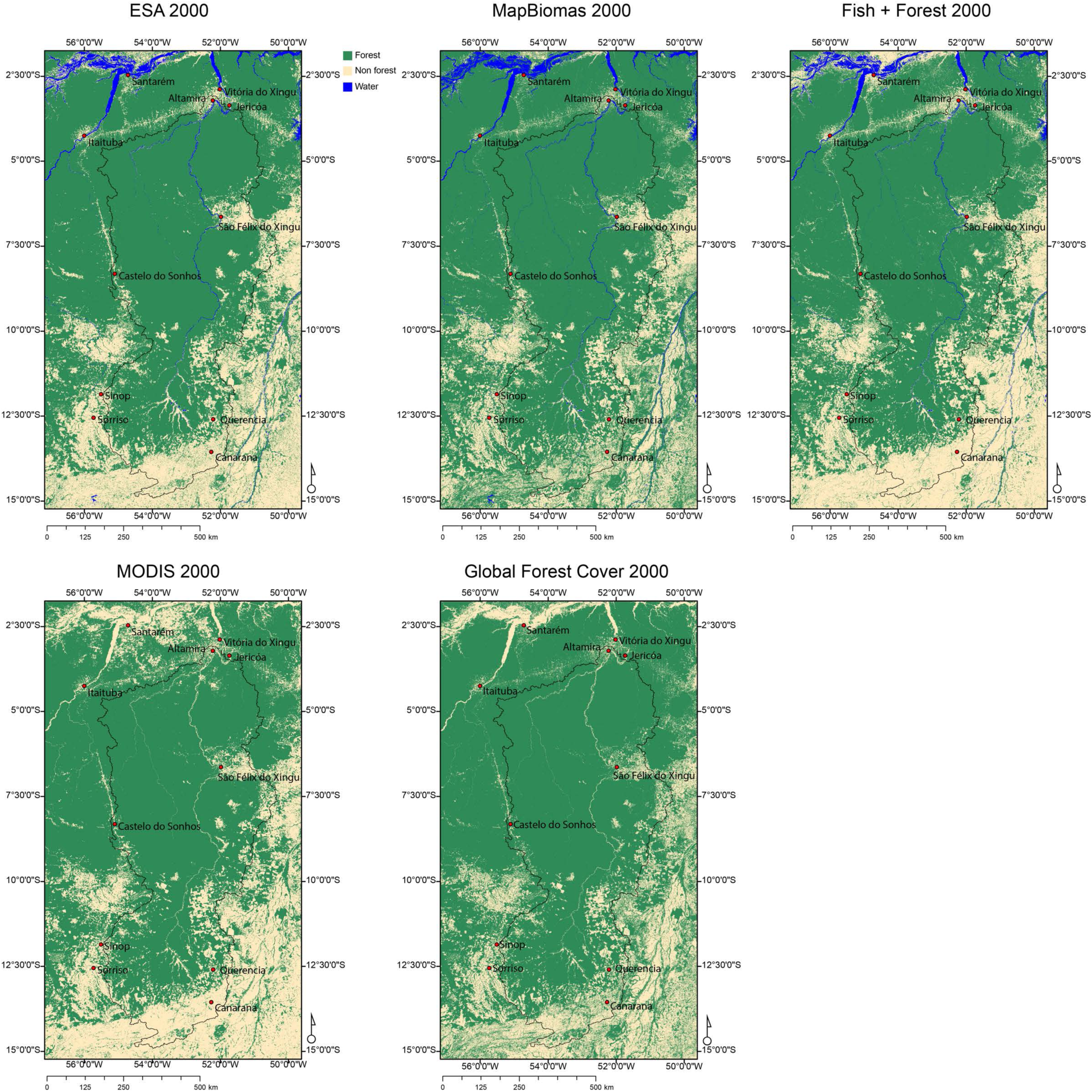
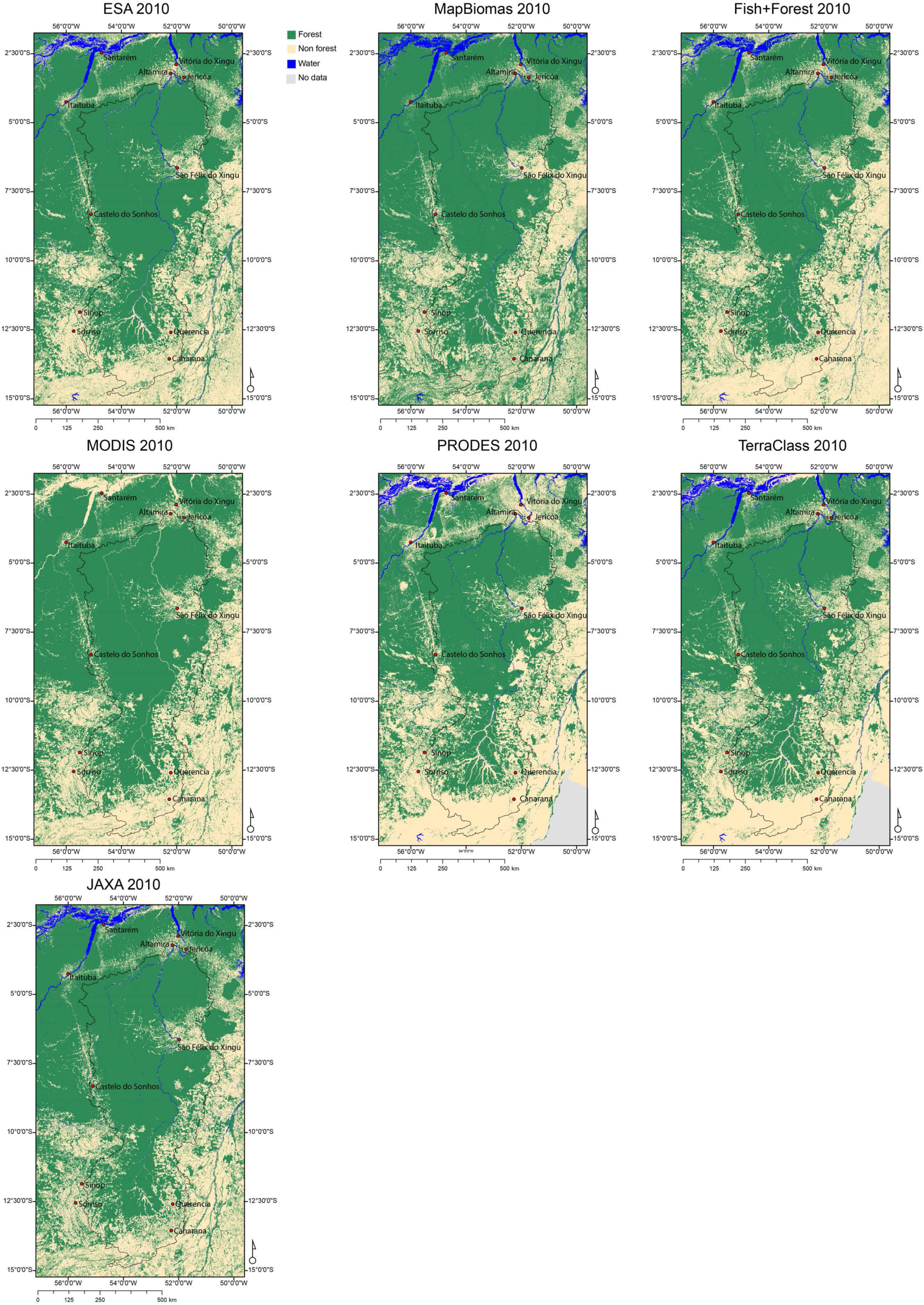
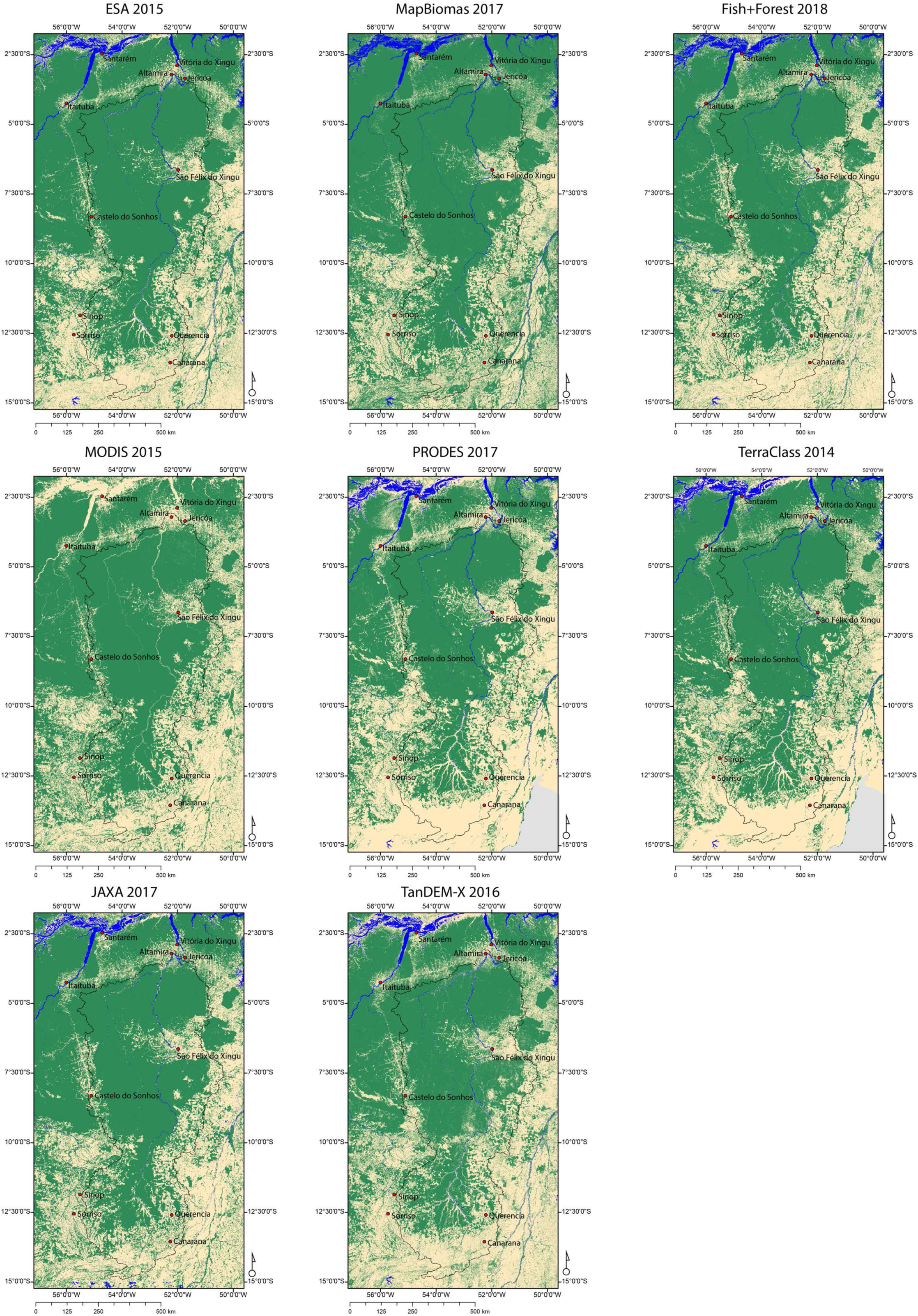
Forest /Non-forest classification datasets examined for the periods A) 1989-1992, B) 2000, C) 2010, D) 2014-2018. For PRODES and TerraClass the State of Goiás is greyed out because the results are for Amazonia only. Locations of cities and the Jericoá rapids are shown for spatial reference.

At such a large scale (Figure 4), most of the datasets appear similar and as such they may seem interchangeable. However, the forest class statistics reveal differences in the total area classified as forest (CA) (Figure 5). For example, for the year 2000, the difference between the minimum (67% Fish + Forest) and the maximum (76.2% Global Forest Cover and MapBiomas) is just over 120,000 km^2^ (12M ha). For the most recent time period, that difference increases to just over 220,000 km^2^ (22M ha) between PRODES (52.5%) and MapBiomas (69.3%). It is important to note that the portion of the State of Goías in the study area (3% of total area) is not included in the PRODES estimates. The deforestation trend between datasets is largely consistent (Figure 5); there is a clear decrease in forest area over the four time periods. While most datasets continue to show a decrease in forest area between the 2010 and most recent period (2014-2018), the MapBiomas and PRODES datasets indicate a slight increase in forest area. For PRODES the reason is likely due to cloud cover contamination in the 2010 data (also seen in Figure 4c) leading to a lower overall area of forest. The JAXA data indicate the greatest forest loss between those two time periods.

**Figure 5.**
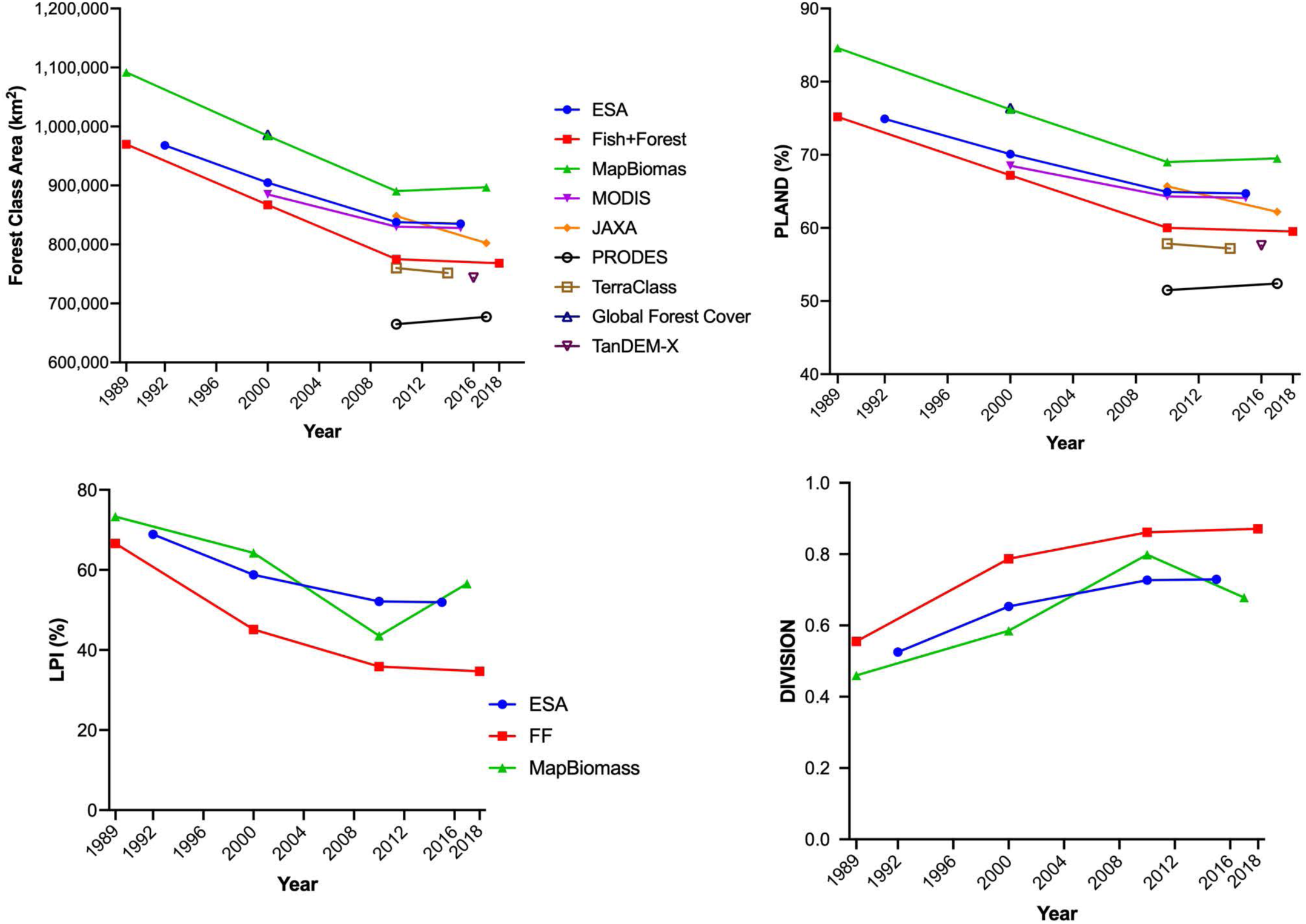
Comparison of total forest area change and fragmentation metrics for the land cover classifications. A) Total forest area (km2), B) PLAND, C) LPI, D) DIVISION. Metrics of PLAND, LPI and DIVISION are only shown for the datasets available for all four time periods.

The PLAND trends (Figure 5) are consistent with CA, with forest loss ranging from 10.3% between for ESA, to 15.6% and 15.1% for Fish + Forest and MapBiomas, respectively. There is also an overall decrease in the area of the largest patch (LPI) over the four time periods for ESA and Fish + Forest (Figure 5c) consistent with temporal erosion of the main core of the intact forest (Figure 6a). In contrast, the LPI indicates a recovery of the largest patch area from 2010 to 2017 from MapBiomas (Figure 5c). Furthermore, DIVISION shows that a consistent fragmentation process is occurring in this area over the four time periods based on the ESA and Fish + Forest classifications (Figure 5d). As with the LPI metric from MapBiomas, DIVISION also indicates a decrease in fragmentation between 2010 and 2017 for this dataset (Figure 5d).

**Figure 6a.**
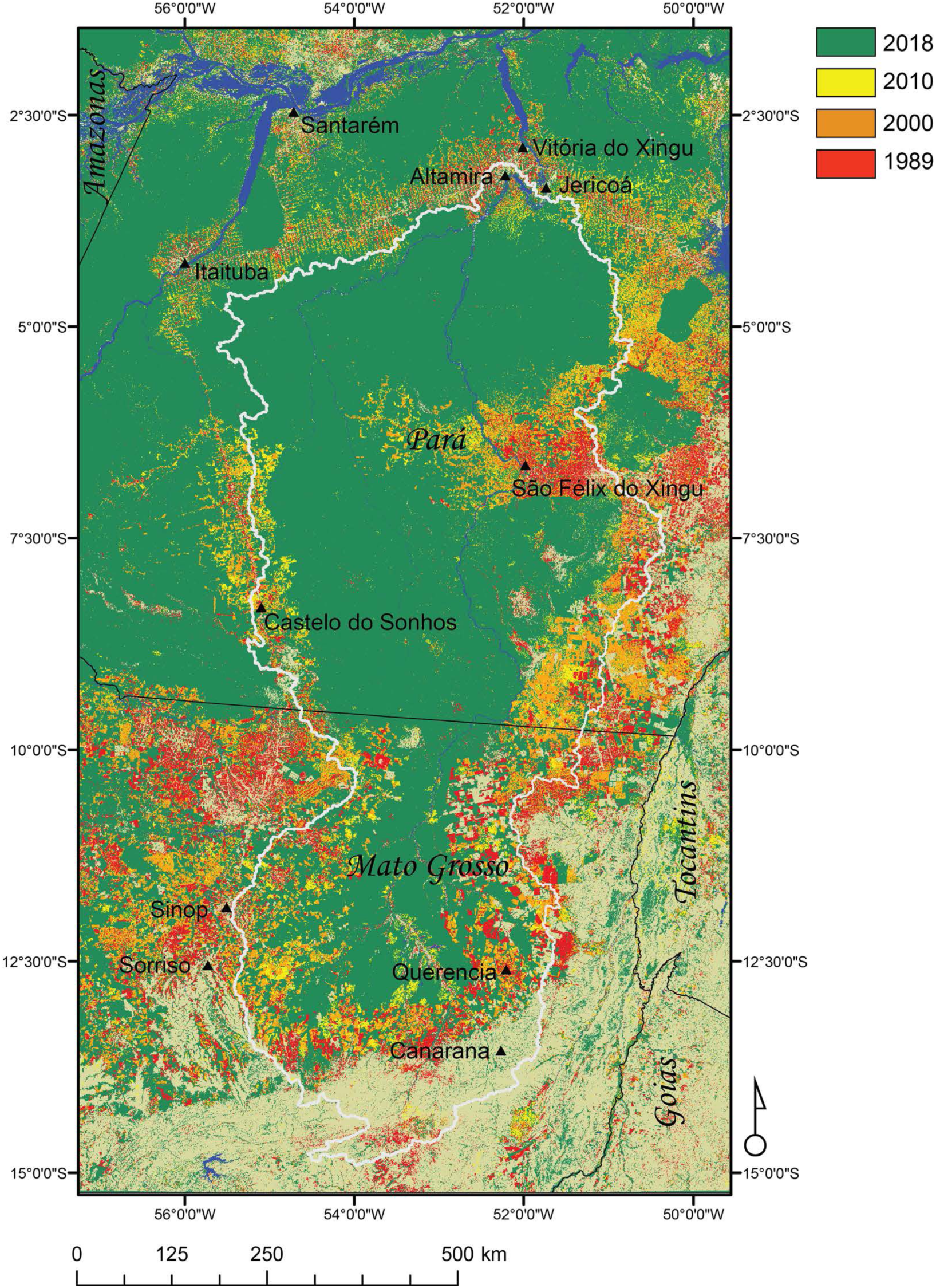
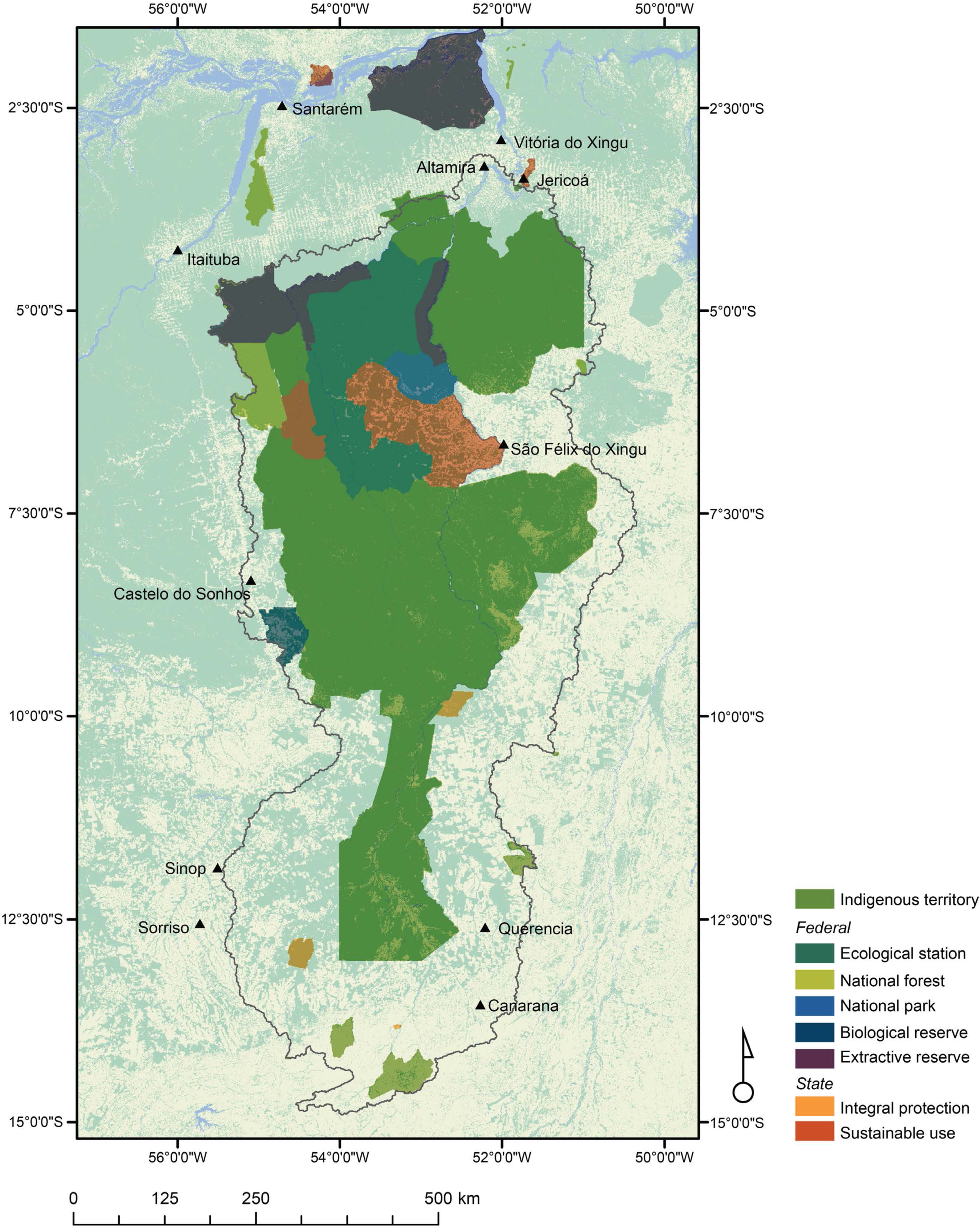

Temporal visualization of deforestation (Figure 6a) illustrates a gradual reduction of the forest cover with the deforestation encroaching upon the core forest areas. While substantial deforestation occurs in the state of Mato Grosso, in Pará the effect of roads (e.g. north of Castelo do Sonhos and between Itaituba and Altamira) leading to increased deforestation is also evident. Marked deforestation for urban growth and increased agricultural development around São Felix do Xingu can be seen between 1989 and 2000 with forest loss in the sustainable use area (Figure 6b) west of the town. The majority of the remaining intact forest in both Pará and Mato Grosso is within the boundaries of protected areas and Indigenous territories. Heavy deforestation is also seen in the Volta Grande and around Altamira after 2010, attributable to construction of the Belo Monte dam complex (Da Silva et al. 2018b) (Figures 1, 6a).

The spatial FAD (Figure 7) and entropy (Figure 8) results illustrate large differences in the landscape (i.e. fragmentation, connectivity and intact forest) depending on the choice of forest cover map. For the oldest time period (Figure 7a), the most noticeable difference is the lack of intact forest west of Querencia from the ESA data. Due to the coarser spatial resolution of the ESA data (300 m vs 30 m for MapBiomas and Fish+Forest) the areas of forest in Mato Grosso at this time are not large enough to be considered ‘intact’ based on the multiple scales of assessment in the FAD metric (i.e., the largest scale of FAD is 243 pixels which results in a moving window of 72.9 × 72.9 km for ESA). There is less intact forest around São Félix do Xingu based on the Fish +Forest dataset due to the increased number of small deforestation patches which are not present in the MapBiomas data. Similar trends can be seen in subsequent time periods with the coarsest spatial resolution data (i.e., EAS and MODIS) showing the least amount of intact forest. For 2000 (Figure 7b), there are considerable differences in the Volta Grande region (Altamira, Jericoá, Vitória do Xingu) between ESA and MODIS. The forest was classified as transitional and dominant between Altamira and Santarém from the MODIS data whereas there is a large patch of intact forest in the same region from the ESA data. From the finer spatial resolution maps, the Global Forest Cover shows the largest continuous patches of intact forest in comparison to MapBiomas or Fish + Forest which indicate a higher proportion of ‘interior’ forest and greater fragmentation of the intact forest class. Interestingly, in 2010 (Figure 7c), the high resolution JAXA dataset (25 m spatial resolution) has large patches of intact forest, similar to the coarse spatial resolution ESA and MODIS data. In contrast, the other 30 m datasets (PRODES, Fish + Forest, MapBiomas, TerraClass) all show a high degree of fragmentation especially west of Sinop and Sorriso, but also throughout Mato Grosso. The same pattern is seen for the most recent time period (Figure 7d) with the exception of TanDEM-X which has diagonal striping artifacts throughout the study area resulting in an erroneous FAD classification. The summary of the area within each FAD class (Figure 7e) shows the decrease in intact forest with a corresponding increase in ‘interior’ and ‘dominant’ classes. For the three most recent time periods variations of >10% of the total area can be seen for the ‘intact’ and ‘interior’ classes between datasets.

**Figure 7.**
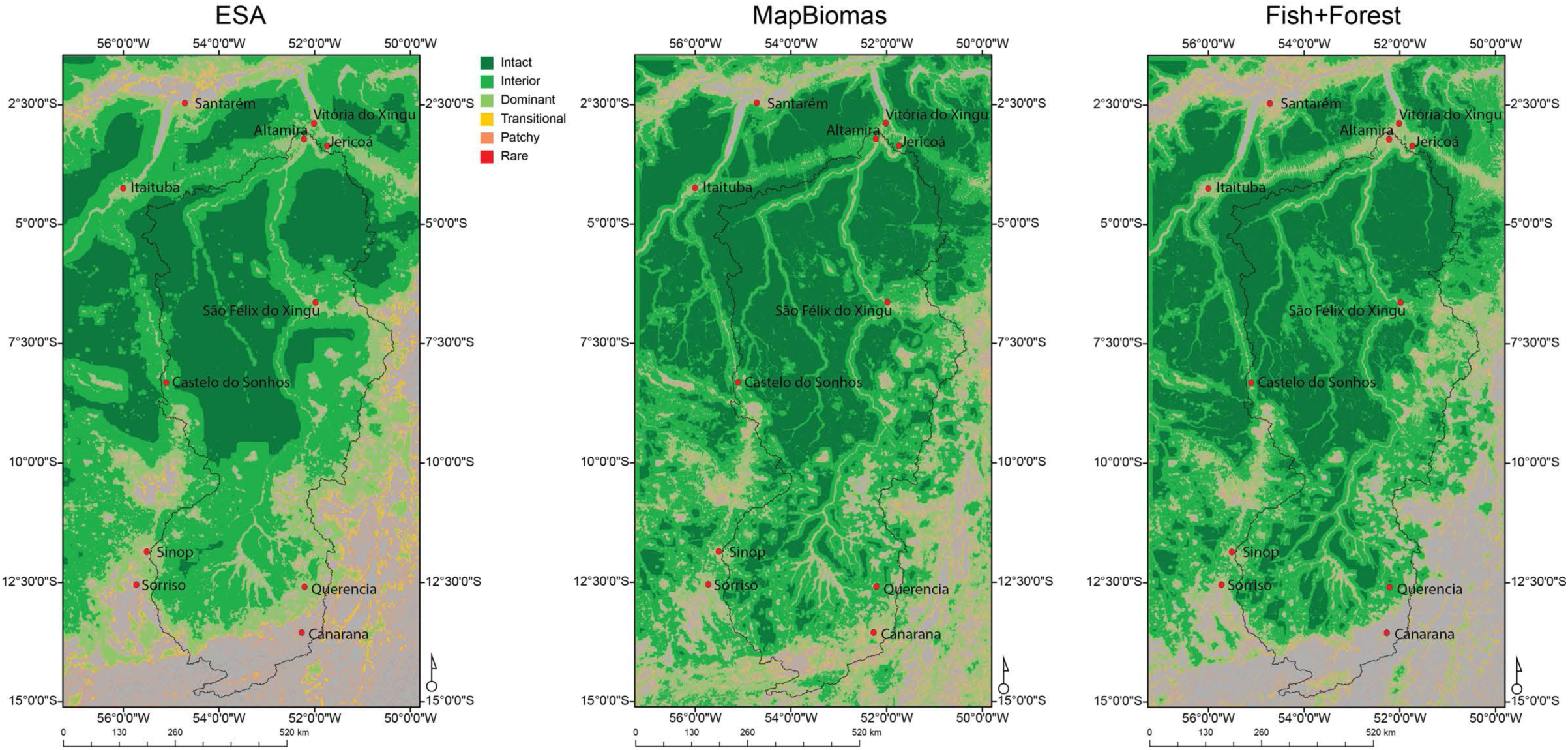
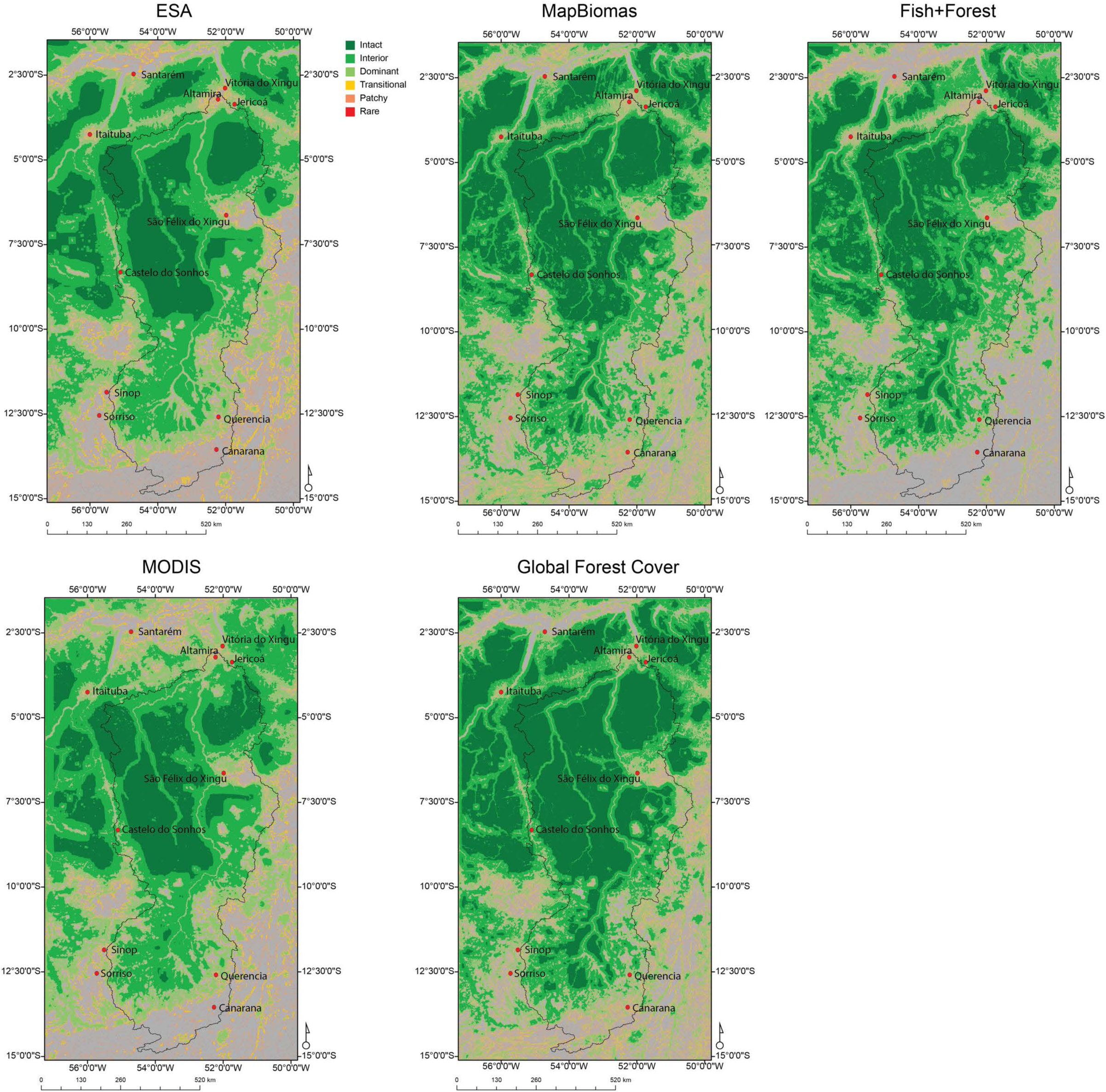
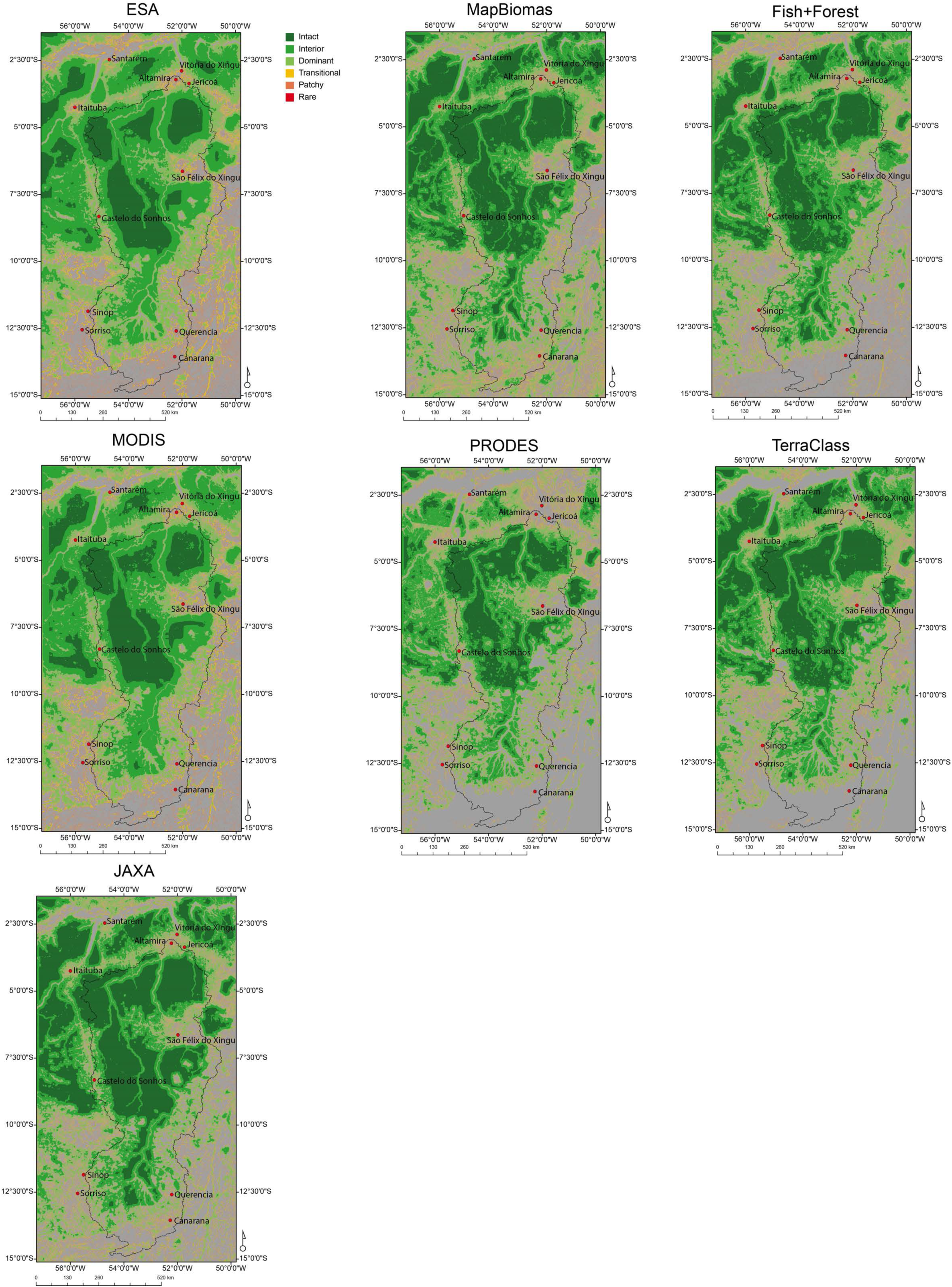
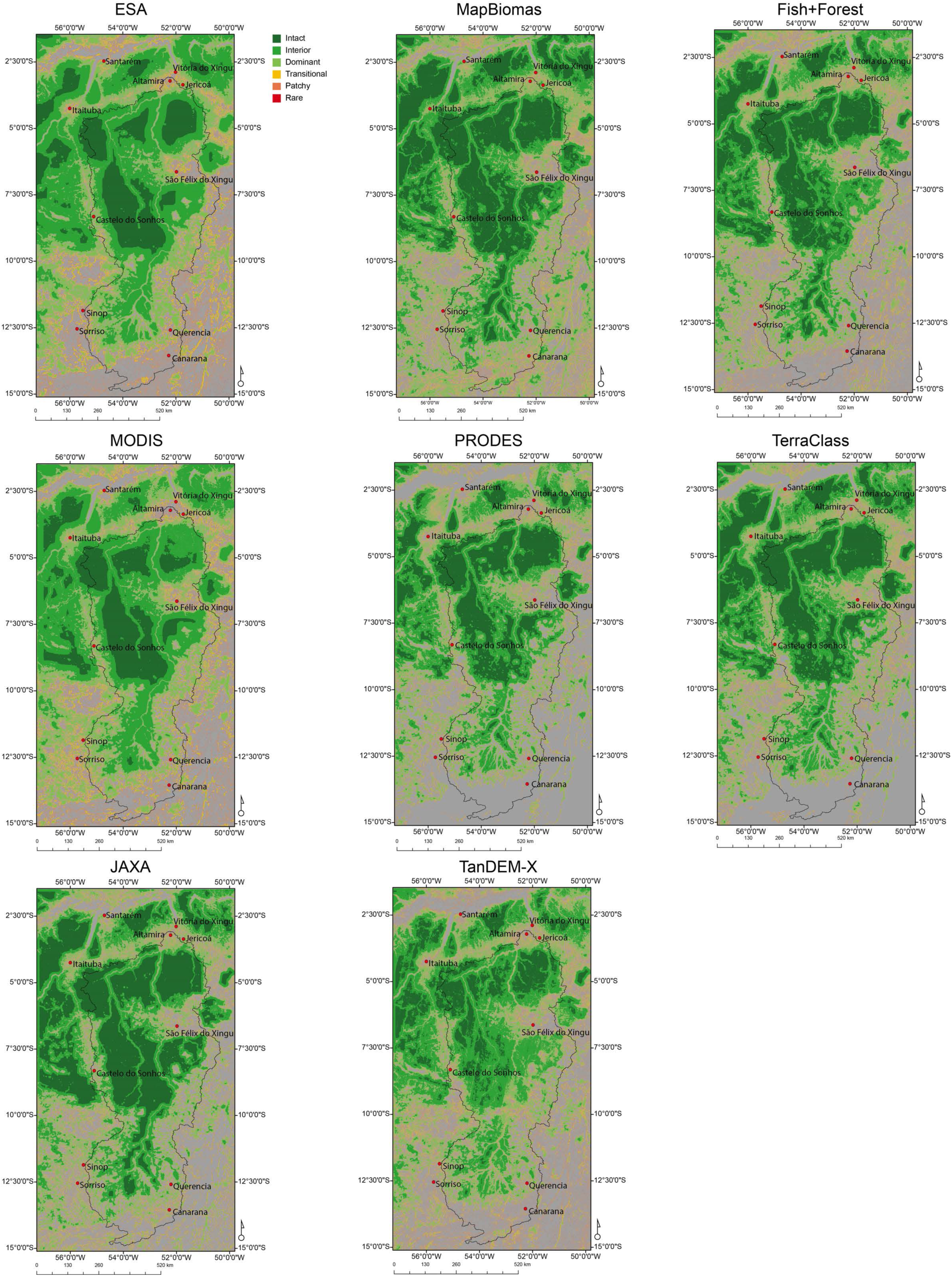
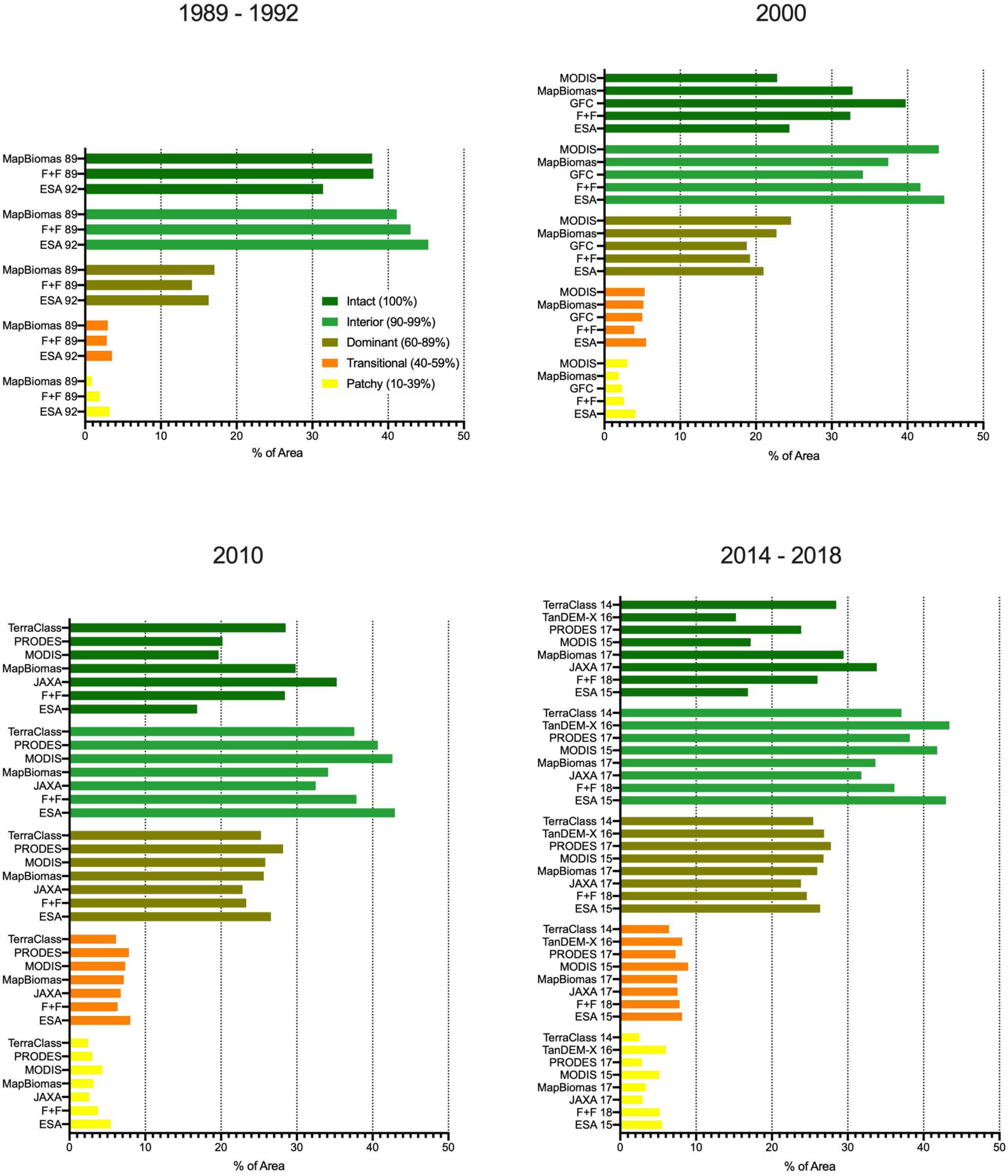
Average Foreground Area Density (FAD) classes for the A) 1989-1992, B) 2000, C) 2010, D) 2014-2018 time periods. E) Percent area of each FAD class from the classifications.

**Figure 8.**
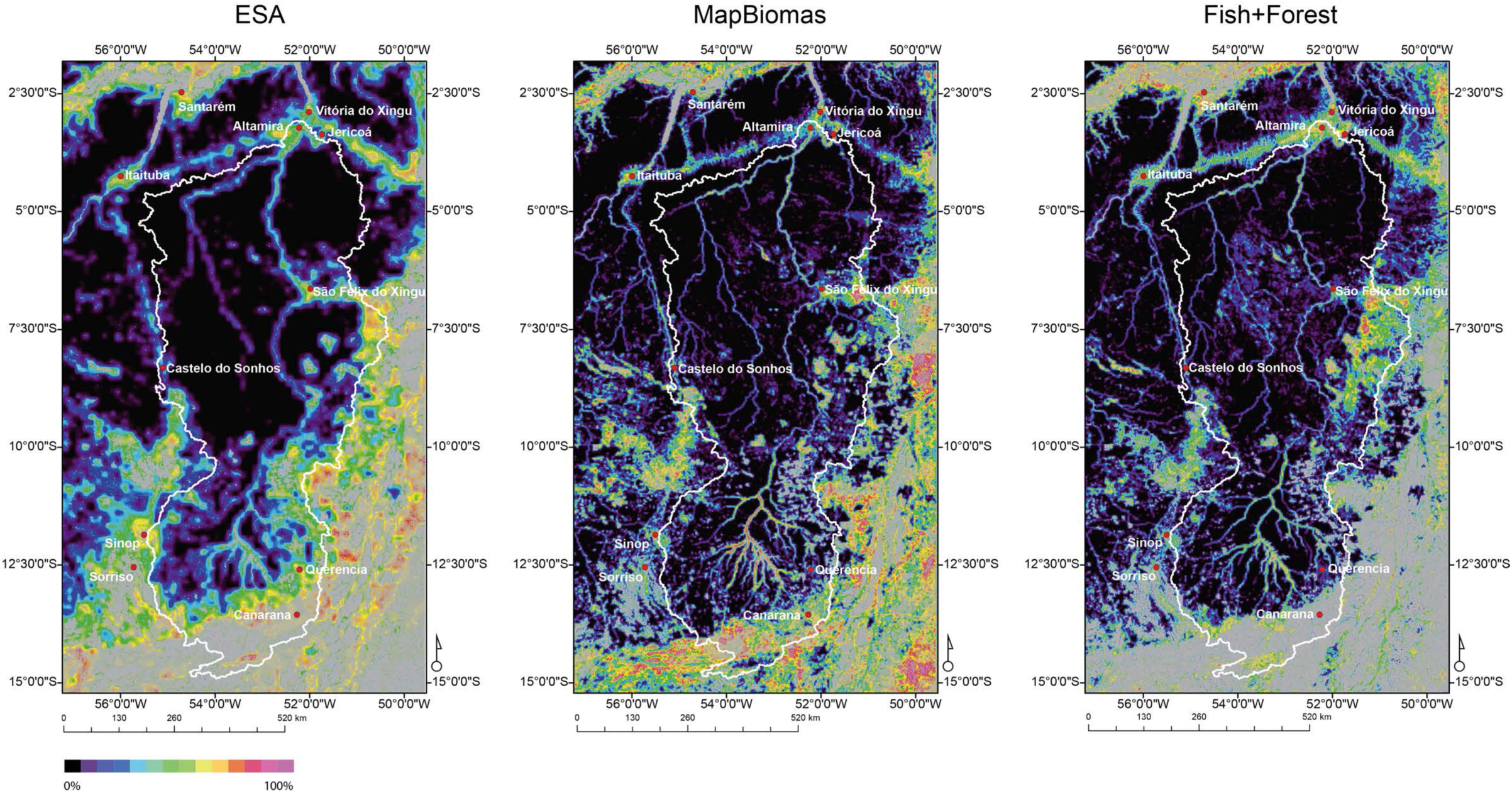
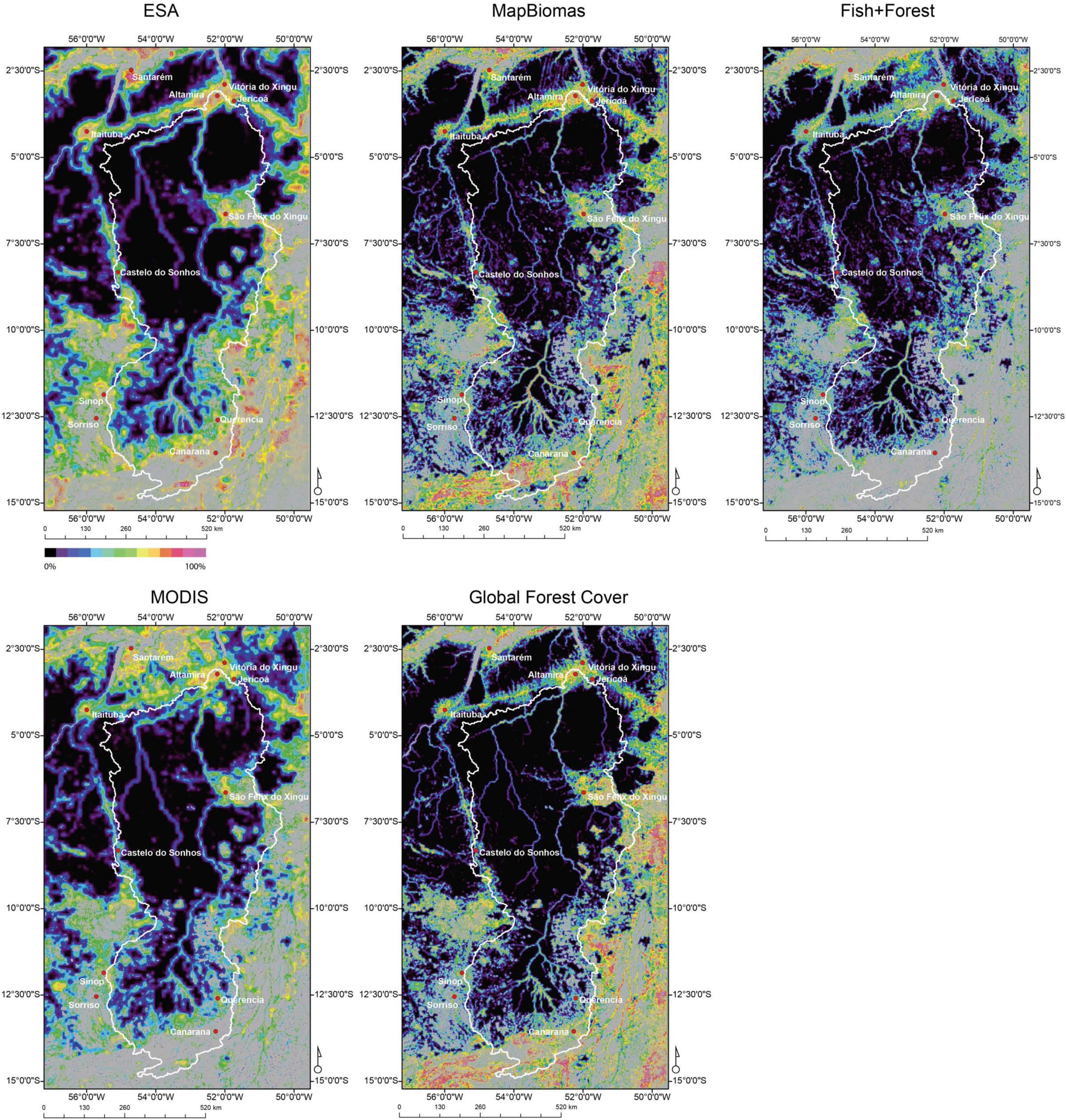
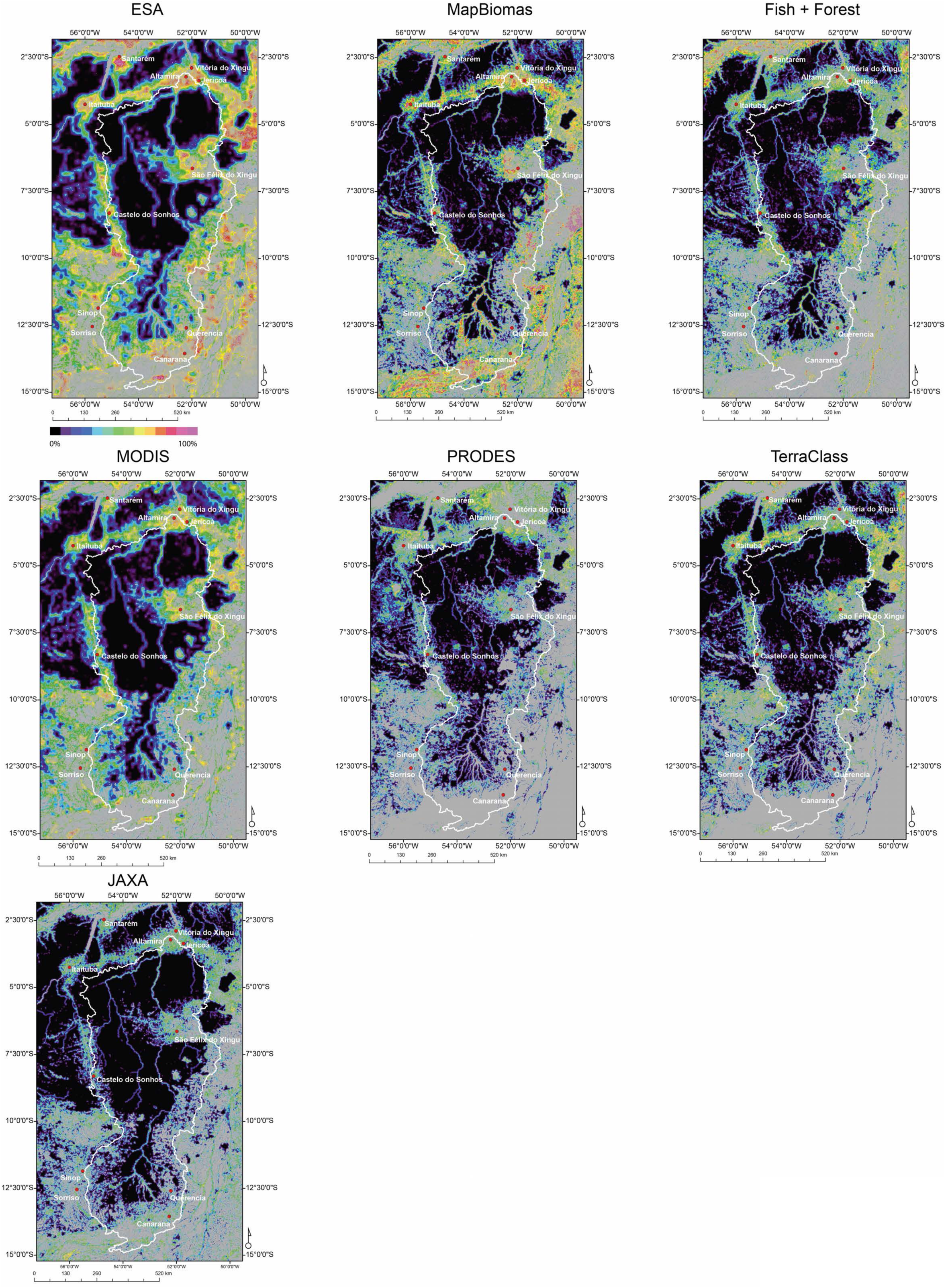
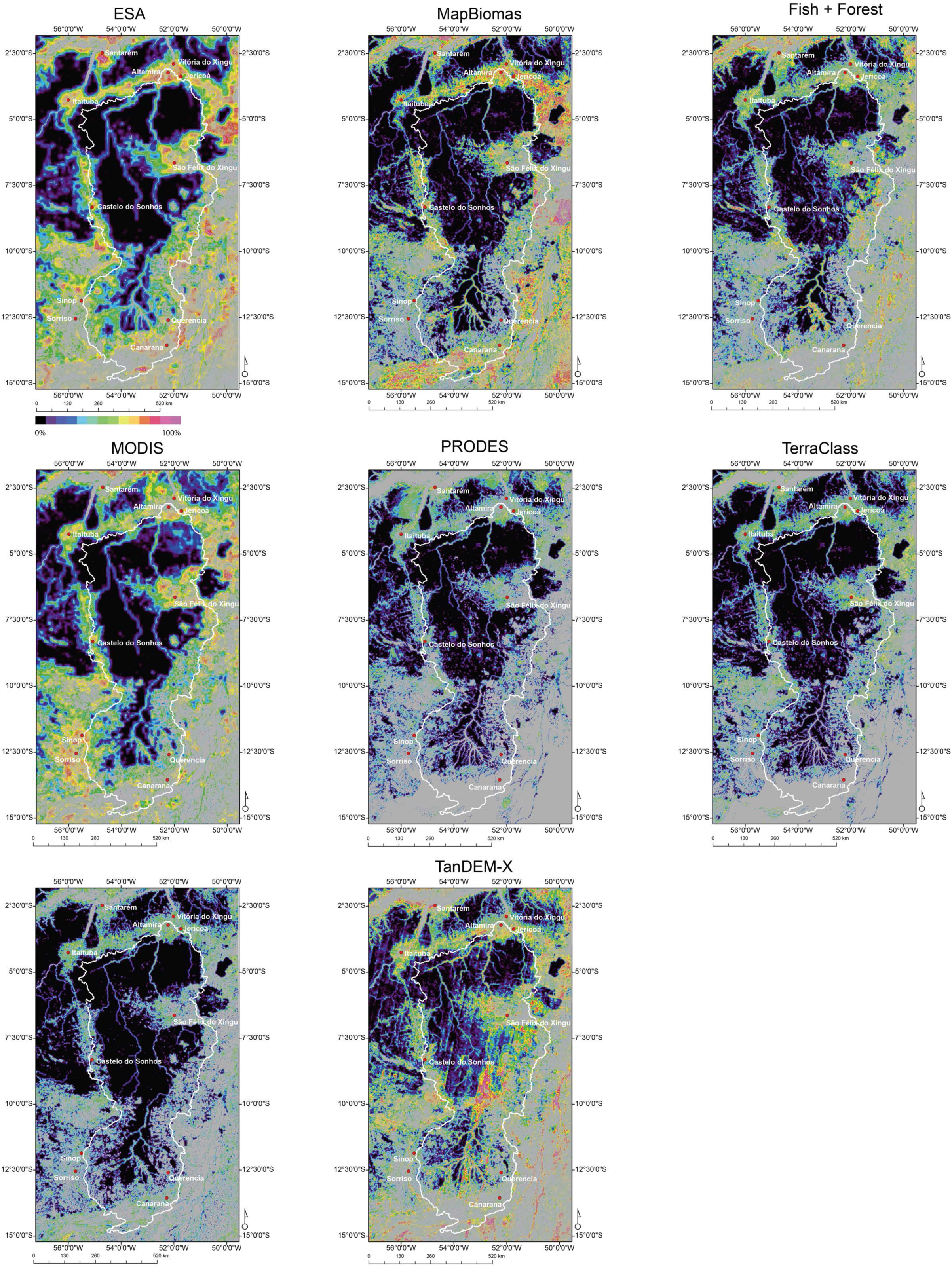
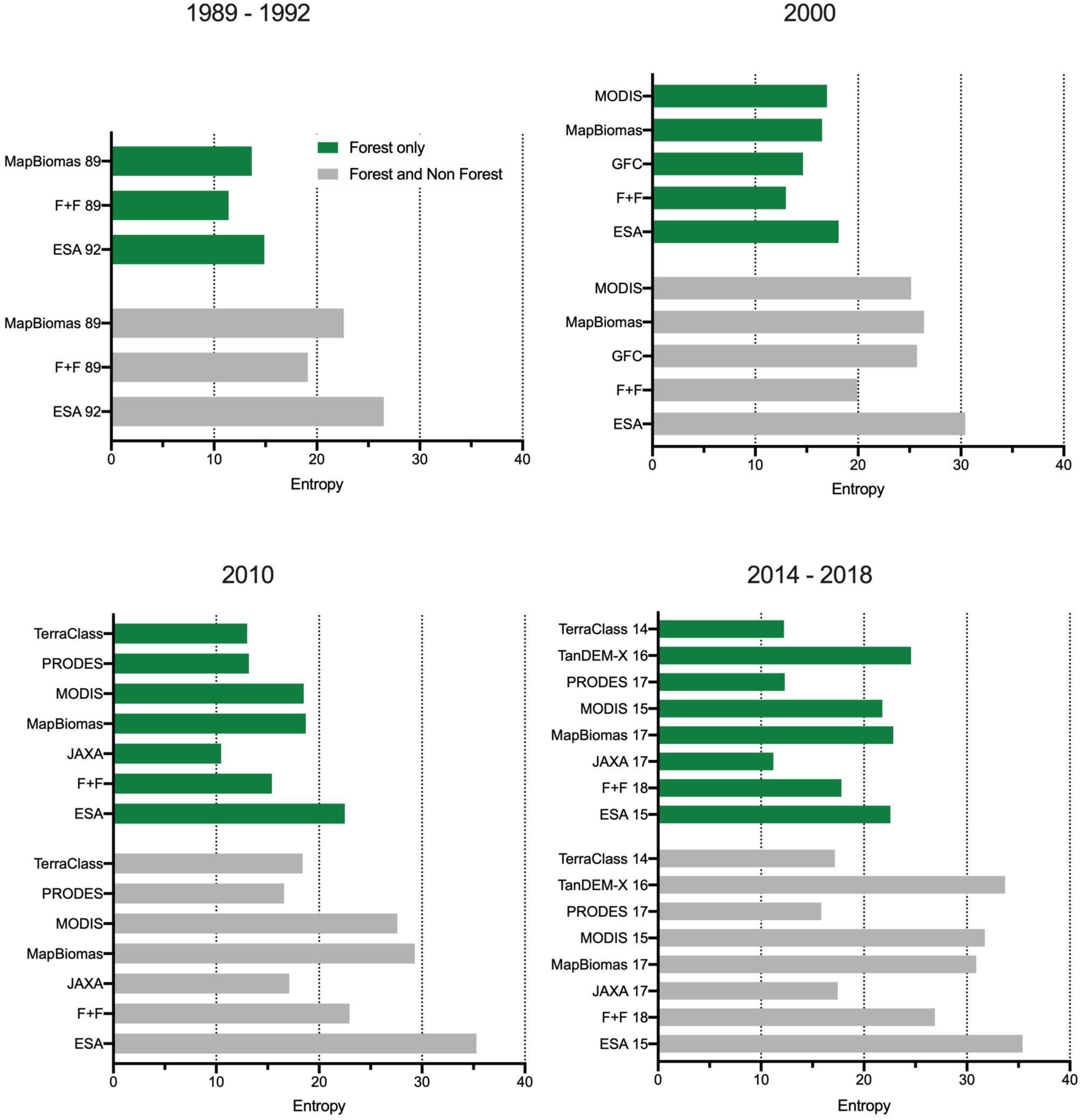
Spatial entropy for the A) 1989-1992, B) 2000, C) 2010, D) 2014-2018 time periods. E) Overall measure of entropy for the forest and non-forest classes from each classification.

The entropy metrics (Figure 8) illustrate the same general pattern as the FAD (Figure 7), but the differences between datasets are more pronounced. For example, even in the oldest time period (Figure 8a), substantial differences between the three datasets can be seen, especially in the north between Itaituba and Altamira with MapBiomas indicating the greatest amount of fragmentation. For the other two data sets, ESA and Fish + Forest, increased entropy can be seen along the road between Itaituba and Altamira, but the fragmentation does not extend as far into the surrounding forest. For the year 2000 (Figure 8b), very high degrees of fragmentation can be seen from all datasets in Mato Grosso and Goías. The larger number of small forest patches in those states in the Global Forest Cover and MapBiomas datasets indicates the highest amount of fragmentation. For 2010 (Figure 8c), ESA and MapBiomas have the largest number of small dispersed patches of forest in the landscape leading to the most fragmented landscape outside of the main intact forest region. That same pattern can be seen for the most recent time period (Figure 8d) for ESA, MODIS, MapBiomas and TanDEM-X. However, the striping artifacts in TanDEM-X greatly increase the perceived fragmentation in the landscape. The summary of the overall entropy values (Figure 8e) reiterate an increase in fragmentation over time. Of note is the relatively low value of entropy for the forest class from the JAXA dataset for 2010 which indicates a less fragmented forest than any of the datasets from the 1989-1992 period. This is likely due to the dataset missing small deforestation patches because of its low forest cover threshold (10%) in comparison to the others which use a 30-60% threshold.

The scatterplot between core (i.e. interior) and edge (i.e. perimeter) from the MSPA statistics further reveal the differences in the spatial patterns of forest fragments in the study area over time between the various datasets (Figure 9). Overall, an expected negative correlation indicating an increase in edge with decrease in core area can be seen. However, the placement in those two axes of the individual datasets representing the same time periods varies greatly. As an outlier, the TanDEM-X map from 2016 indicates the lowest core area in comparison to the amount of edge. This, however, is likely an artifact due to the striping seen in Figures 7 and 8. JAXA, TerraClass and PRODES show the least amount of attrition in core area over time compared to the other datasets, whereas ESA has the least amount of change between the two most recent time periods.

**Figure 9.**
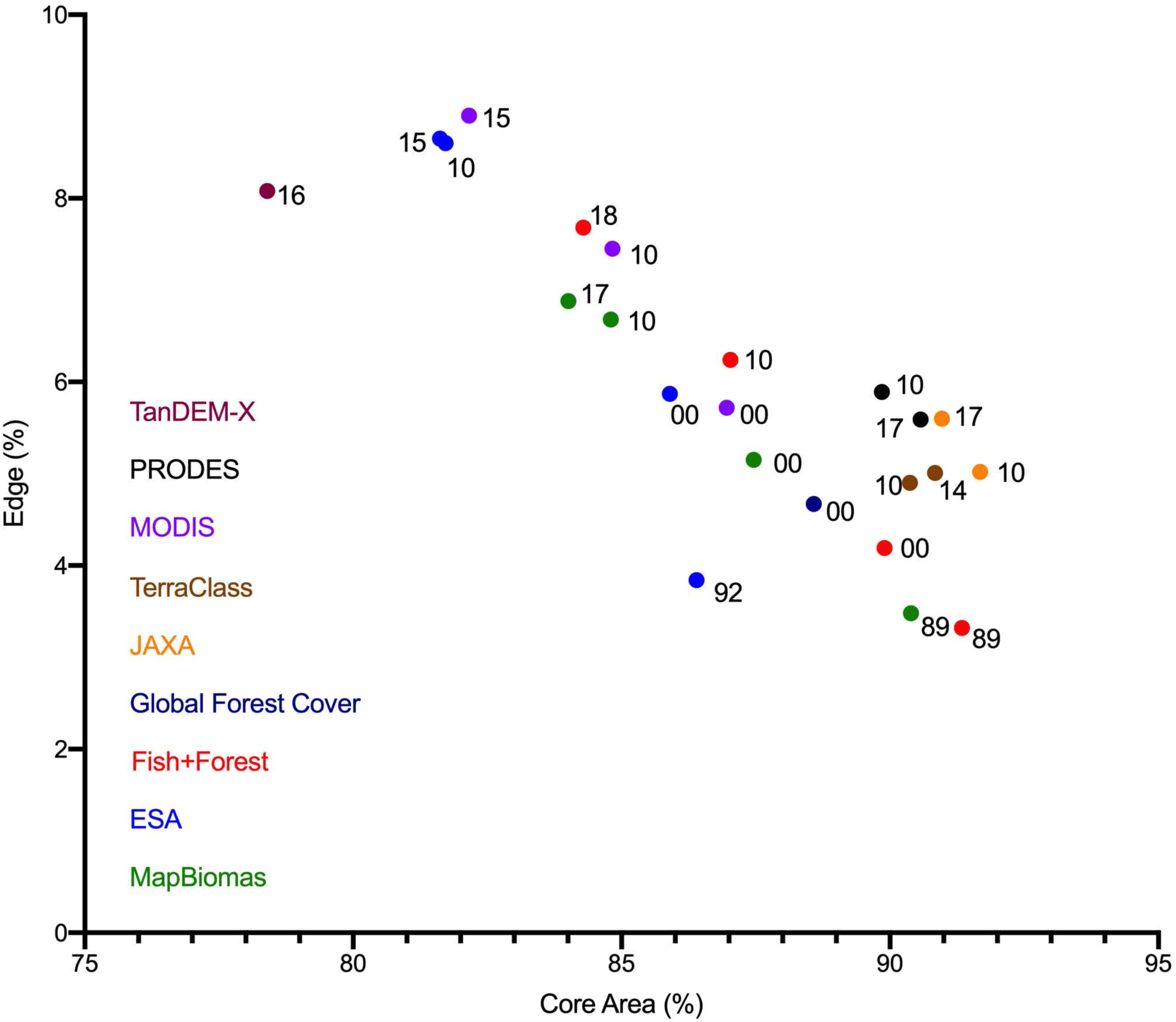
Relationship between MSPA statistics ‘core area’ and ‘edge’ for the forest class from the different datasets. The numbers represent the year of the classification.

Of the water layers examined here, only the EC JRC/Google and HydroSHEDS are explicitly hydrological versus water being another land cover class extracted as part of a land cover classification (Figure 10). Over the entire study area, from the most recent time period, a large range in total surface water extents can be seen (Figure 10a). From the six maps dated after Belo Monte was operational, the area of surface water ranges from 22,790 km^2^ (TanDEM-X) to 44,325 km^2^ (EC JRC/Google 1984 – 2018). To avoid misclassification, water was not directly assessed by the TanDEM-X product (50 m pixels) it was added from an ESA CCI land cover data product where it is mapped at 150 m pixel size mainly from radar acquired over the 2005 – 2012 period (Table 2) (Kirches et al. 2014; Matone et al. 2018). In contrast, the EC JRC/Google 1984 – 2018 product is from Landsat time sequences produced at 30 m pixel size. The product represents the maximum extent of the water observed over the 34-year time period. The focus on the Volta Grande (Figure 10b) indicates a lack of spatial consistency in the estimation of surface water between the different datasets. An absence of river connectivity can be seen in the water layer from the ESA data near Jericoá where the natural channels have complex fluvial geomorphology (Figure 11a); the junction of the Xingu with the Iriri river, a large tributary, similarly lacks connectivity in the ESA data. This is expected due to the large pixel size of the ESA data in comparison to the relatively narrow channels in this area of the river. Similarly, the JAXA data shows discontinuity throughout the Xingu river south of Altamira. The TerraClass and PRODES datasets have the largest extent of surface water in this area. Nevertheless, both of these datasets oddly lack the Bacajá river, a southeast tributary entering the Volta Grande stretch of the Xingu (Figure 12), which is present in every other dataset examined here and can also be seen in the satellite image from 2017 (Figure 10b).

**Figure 10.**
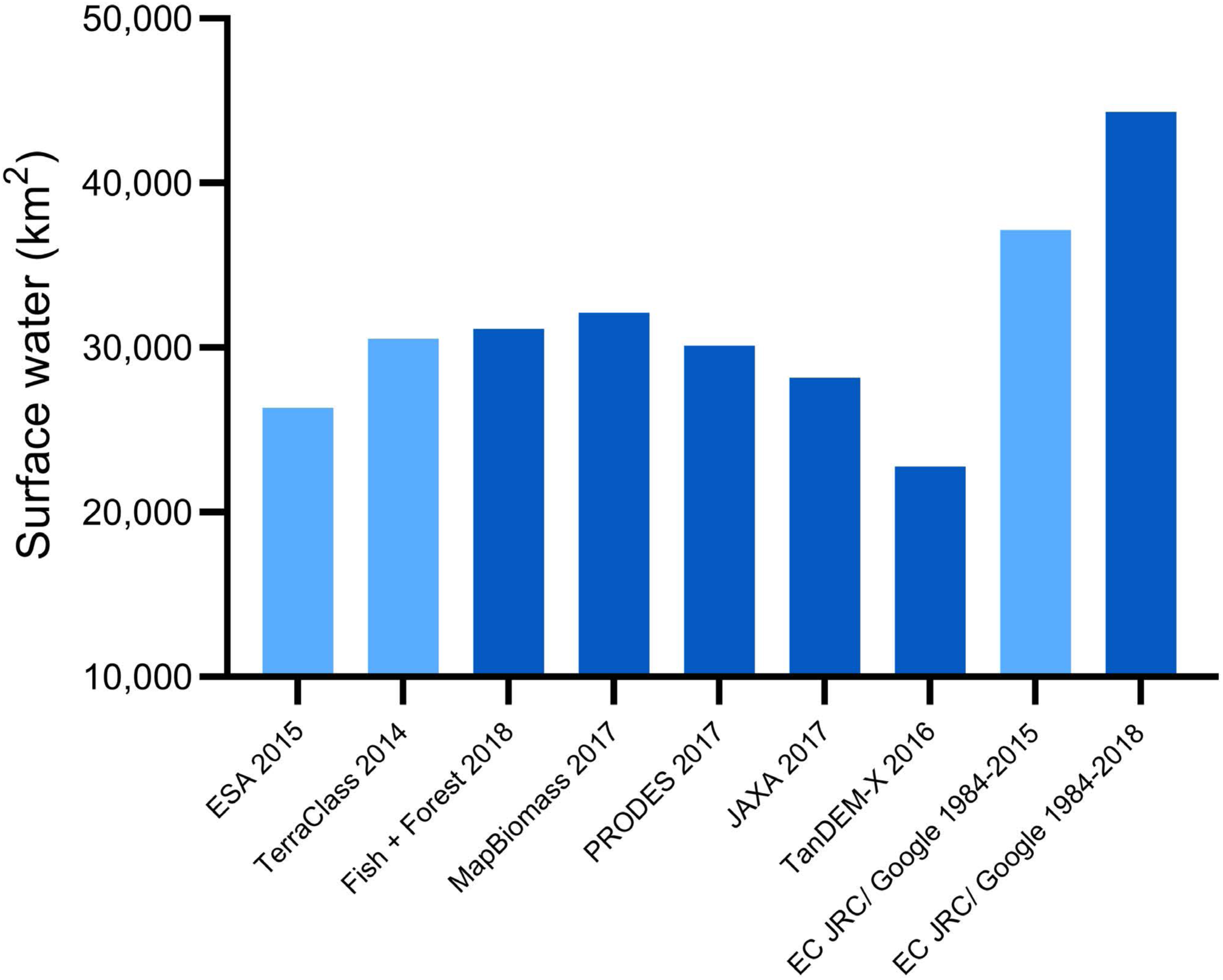
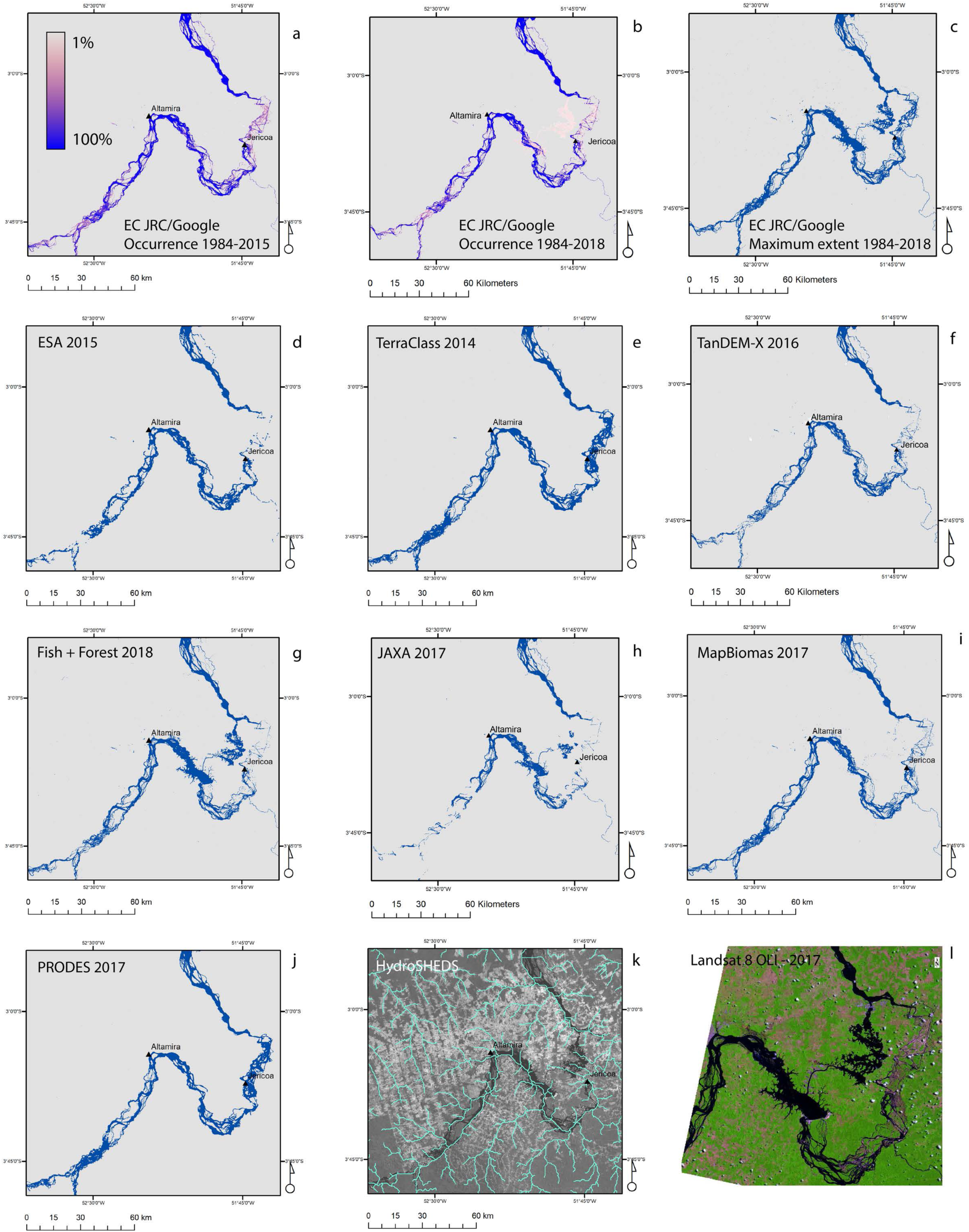
A) Total surface water mapped by the dataset from the most recent time period over the study area. The three light blue columns indicated maps from before the operationalization of Belo Monte. The dark blue columns indicate maps from after the operationalization of Belo Monte. The two EC JRC/Google datasets illustrate the maximum extent of surface water mapped over the given time periods. B) Examples of surface water classifications with the Volta Grande area is shown in detail. a) Probability of surface water occurrence over the 1984-2015 period (EC JRC/Google), b) Probability of surface water occurrence over the 1984-2018 period c) Maximum extent over the 1984-2018 period (EC JRC/Google), d) ESA 2015, e) TerraClass 2014, f) TanDEM-X 2011-2016 period, g) Fish+Forest 2018, h) JAXA 2017, i) MapBiomas 2017, j) PRODES 2017, k) HydroSHEDS overlain on a single band from a Landsat OLI 8 image from 2017 for context, l) Landsat 8 OLI image acquired in 2017 illustrating the Belo Monte reservoir and the dewatered section of the Volta Grande.

**Figure 11.**
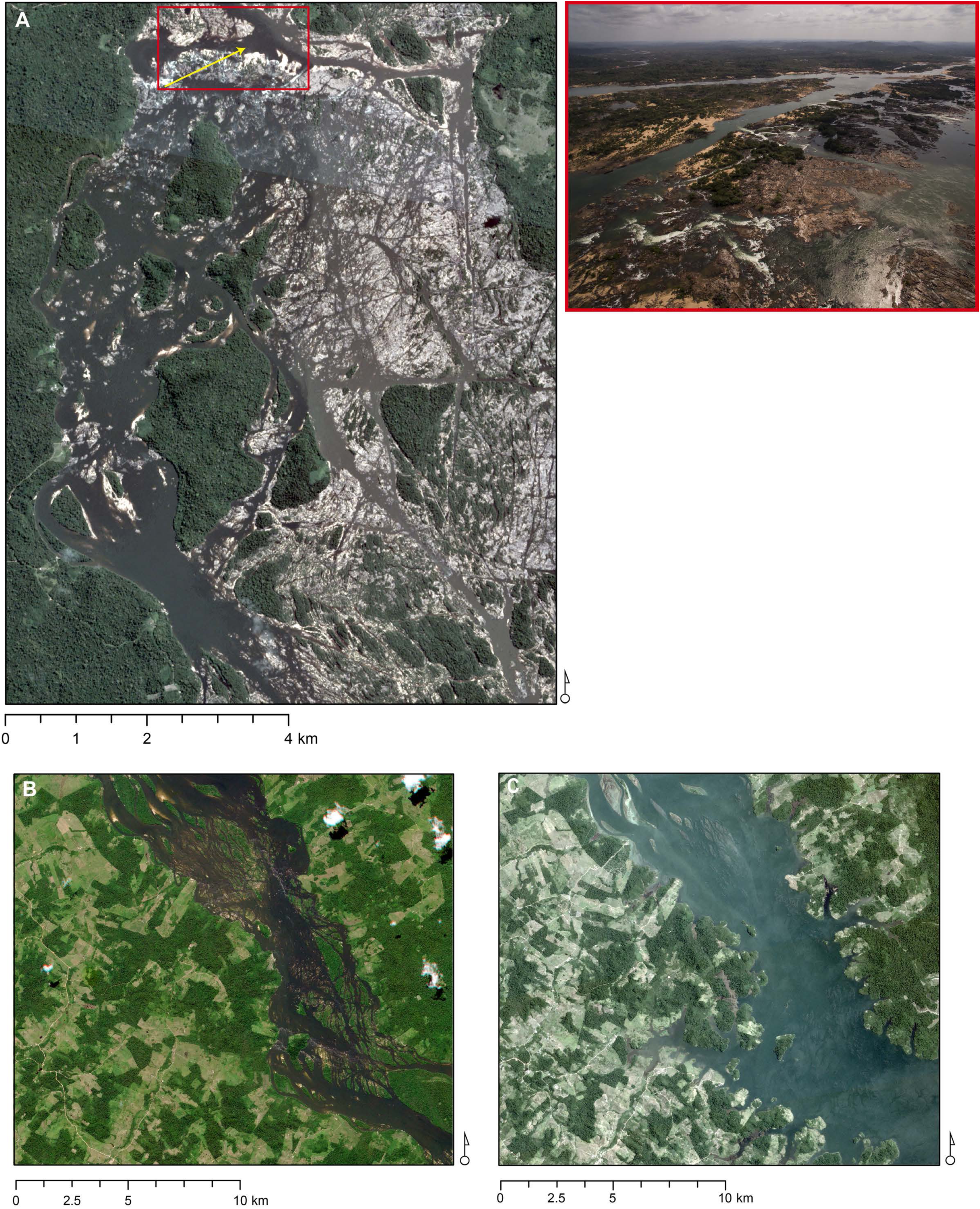
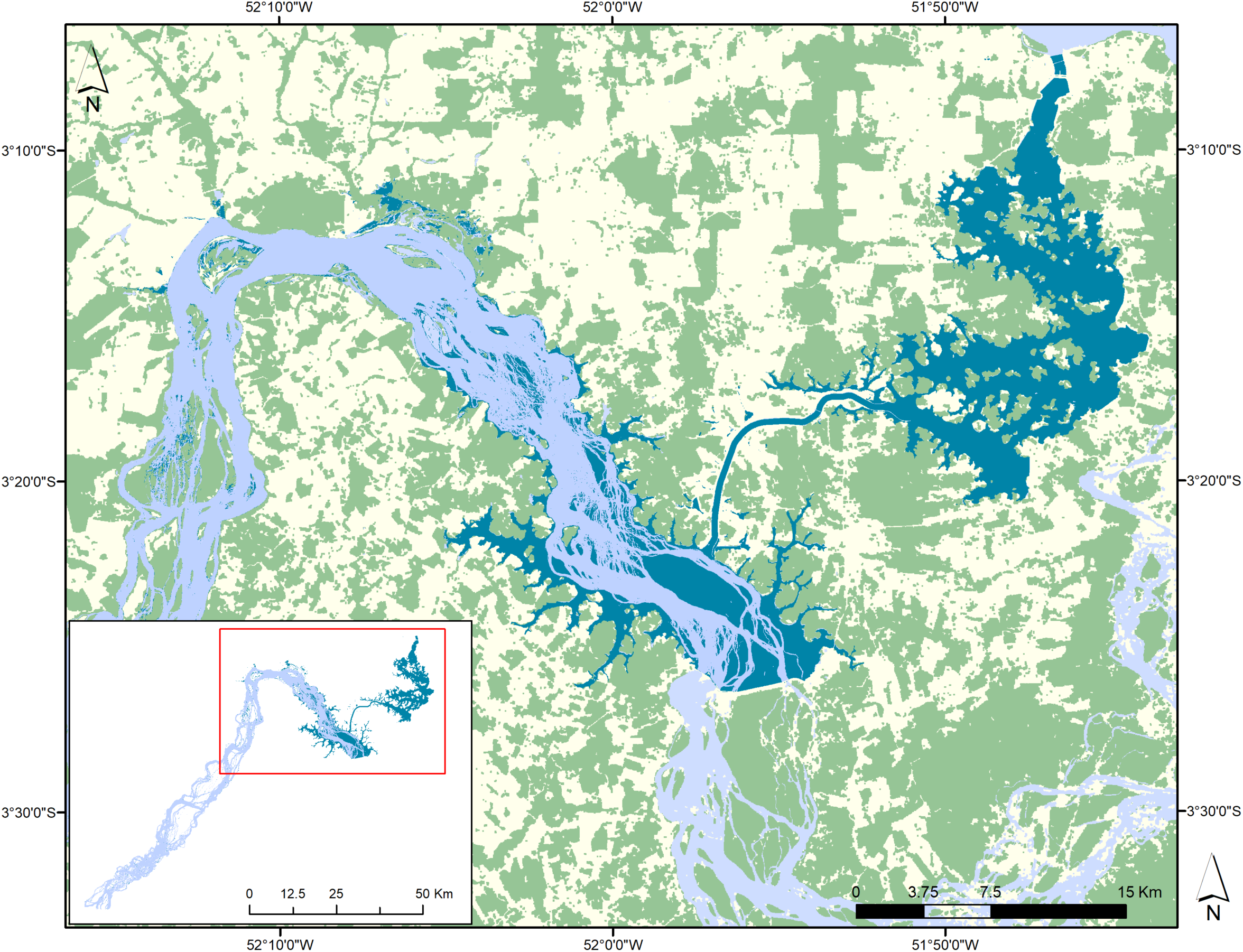
A) Complex fluvial geomorphology of the Xingu river in the dewatered zone. PlanetScope mosaic representing the April – June period 2017 is shown with the Jericoá rapids highlighted by the red rectangle. A UAV photograph from September 2015 of the area in the red rectangle is shown. Yellow arrow indicates the look direction of the UAV photograph. Large areas of exposed rock and sand can be seen in the photograph. B) Example of the original course of the Xingu river between Altamira and the Pimental dam (RapidEye image from 2011). C) Same area as in B following the flooding of the reservoir (PlanetScope image from 2019). D) Map of the extent of the Xingu river from its confluence with the Iriri river and the Pimental dam (Kalacska et al. 2020). The original course of the river is shown in light blue (2011) with the post operationalization of Belo Monte river course and reservoirs shown in dark blue (2019). Data are superimposed on the 2018 Fish+Forest classification.

**Figure 12.**
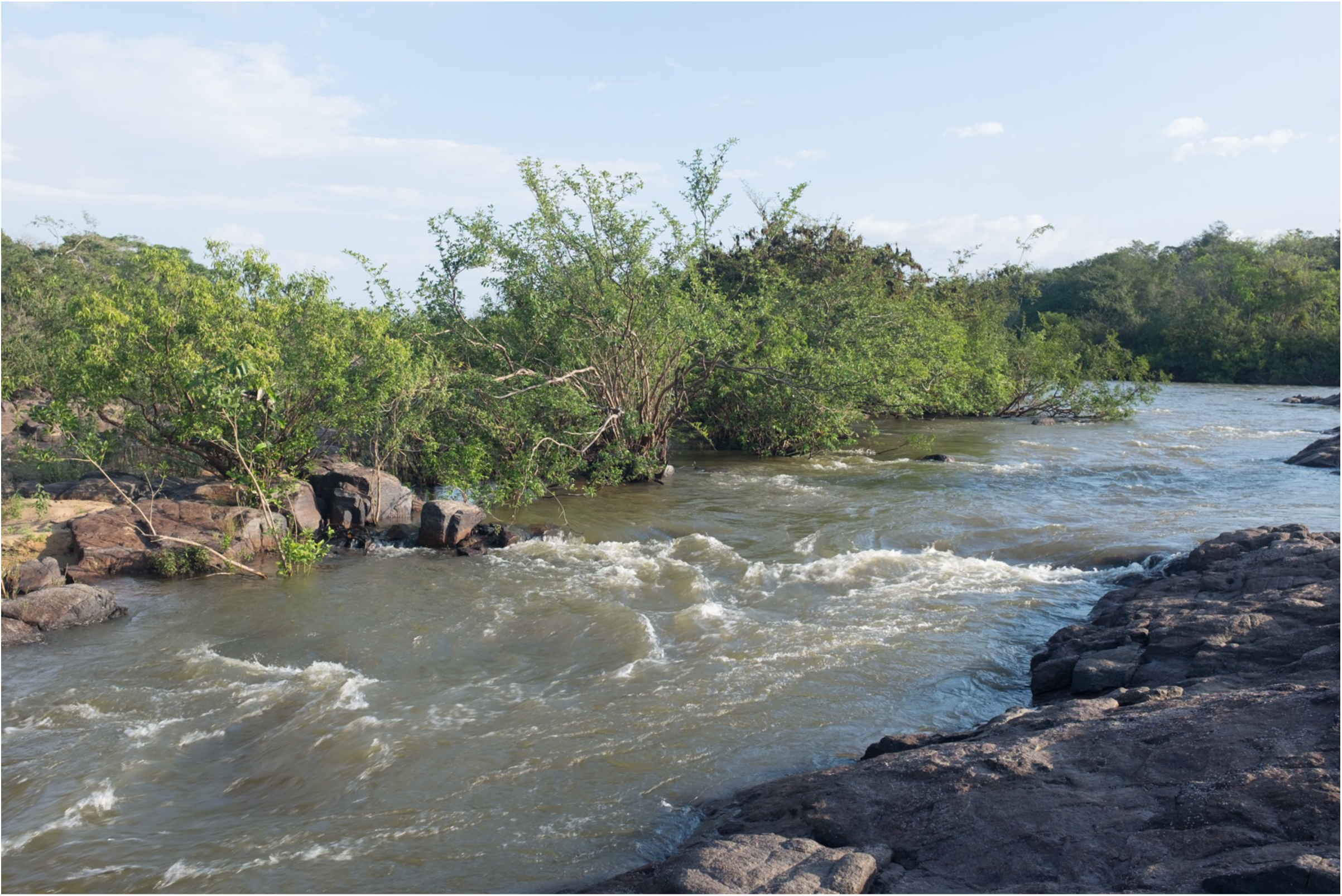
Photograph of the Bacajá river taken at its confluence with the Xingu river in the dry season. As a larger tributary of the Xingu river it should be present in hydrographic layers.

The Belo Monte dam complex became operational in November 2015. Therefore, its reservoir should be present in datasets representing 2016 onwards. In the Landsat image from 2017 (Figure 10b), the flooded channel and reservoir can be clearly seen as well in Figure 2b from July 2016 and the map in Figure 11c. Its absence from the TanDEM-X data (Figure 10b) can be explained by the use of the ESA CCI water layer produced from radar scenes acquired prior to 2015, and the multiple year reference period (2011-2016) over which the radar scenes were gathered. The lack of this large body of water from the PRODES, MapBiomas and TerraClass data sets is more difficult to explain. However, if a pre-existing water layer (i.e., created prior to the flooding) was used without explicit classification of water in the more recent imagery, that may account for the lack of the reservoir in these datasets.

A large difference between observed surface water and modelled river can be seen with the HydroSHEDS dataset (Figure 10b). The modelled river network in this area does not always correspond to the observed surface water, especially in the Volta Grande where the Xingu river is shallow with a complex geomorphology and traverses relatively flat terrain. Similar to the findings of da Costa et al. (2019), the course of the Xingu river from a DEM derived source greatly differs from datasets where the surface water is directly observed, demonstrating a point of failure in this type of terrain for DEM-based surface water models.

## DISCUSSION

Overall, our findings demonstrate the importance of transparency in the generation of remotely-sensed datasets. While it is the producers’ responsibility to document not only the accuracy of a given classification/dataset but also the input sources and methodology used (e.g. inclusion of reference data sets, resolution, validation data source, etc.), it is the user’s responsibility to familiarize themselves with the characteristics and limitations of each data set they choose. Apart from obvious differences in granularity (i.e. spatial detail afforded by the different pixel sizes) and data quality (i.e. cloud cover, striping artifacts) the primary reason for the large differences in forest cover between the datasets examined here is not a failure of the technology, but due to different interpretations of ‘forest’. Over 800 definitions of ‘forest’ exist today (Sexton et al. 2016). Internationally, for example, FAO uses > 10% tree cover while the International Geosphere-Biosphere Program uses > 60%. Individual countries, programs and agencies typically use a definition somewhere in between.

Our results showcase the importance of further considering the spatial arrangement of forest, not only the overall area covered. As shown in the fragmentation analyses, maps may have similar trends, but the total areas of forest and the spatial arrangement of the patches in the landscape may change the outcome of conservation decisions depending on the forest cover map chosen. Users need to take into consideration the appropriateness of the map for their proposed application, not only the popularity of a given dataset. Disagreement between MapBiomas and TerraClass has been shown for classes other than forest (Körting et al. 2020) whereas others have criticized the products for not following good practice guidelines in accuracy assessment (Zalles et al. 2019). Others such as Cunningham et al. (2019) critically assess the utility of the GFC dataset with recommendations to use with caution in area that do not meet criteria including high aseasonal rainfall, low relief, and low cropland area. Likewise, Galiatsatos et al. (2020) found that the GFC dataset should not be used for precise estimates of forest cover change in countries with large areas of forest cover and low levels of deforestation. All maps examined here should only be used after critically considering their suitability and limitations for the questions at hand.

As described in Table 2, several of the datasets examined here relied on Google Earth for the acquisition of validation data for the classifications. While the multi-temporal imagery archived through Google Earth allowed for unprecedented virtual exploration of our planet since its initial release in 2001, care still needs to be taken when relying on it as a source of information. Figure 13 illustrates the Volta Grande at two different zoom levels. At the coarse scale (Figure 13a), despite being accessed nearly five years after operationalization of Belo Monte, the archived imagery does not reflect the massive land cover changes and river alterations. The entire reservoir is missing from the coarse scale view on Google Earth and the ‘2020’ date label can be misleading because based the status of the construction of the Pimental dam, the imagery is from no later than 2014. As seen in Figure 2, by July 2015, construction of Pimental was in an advanced stage and by July 2016 it was operational, and the reservoir had been flooded. At the smaller scale (Figure 13b), the imagery is split between a date of July 2017 with the reservoir visible and August 2010 prior to construction. Google Earth collects imagery from third-parties and also relies on users to request imagery ‘refreshes’, leading to less-popular areas being updated less frequently. Outdated imagery in online basemap services is not unique to Google Earth. Figures 13c-d illustrate the basemap service from the USGS EarthExplorer (showing imagery provided by ESRI), a common platform from which satellite imagery such as Landsat and Sentinel-2 can be downloaded. In this basemap the problem is the reverse as seen in Google Earth with the reservoir accurately displayed at a coarse scale (Figure 13c) but split between outdated and current imagery in higher spatial resolution view (Figure 13d). Despite decades of satellite Earth observation, there are locations where cloud cover is a persistent problem for acquiring cloud free imagery and these two examples showcase the challenges of developing a global cloud free basemap at multiples scales through a mosaic of components drawn from different dates and satellites that have different temporal resolutions. Other than spaceborne video (e.g. VividX2 constellation), geostationary satellites that maintain their position with respect to the rotation of the Earth (e.g. GOES-16), provide the highest temporal resolution (up to 30 sec – 15 min) in order to monitor weather events (e.g. hurricanes), clouds and fires. Their spatial resolution of 500 m to ∼1 km however is too coarse for most freshwater applications (see Figure 3). Daily revisit time, high spatial resolution imagery is currently achieved through constellations rather than individual satellites such as RapidEye which comprises five satellites providing daily imagery off-nadir (5.5 days at nadir) or the constellation of PlanetScope (>130 CubeSats) providing daily multispectral imagery (3.7 m) and SkySat providing up to twice daily multispectral imagery (50 cm). However, imagery from these constellations can be expensive.

**Figure 13.**
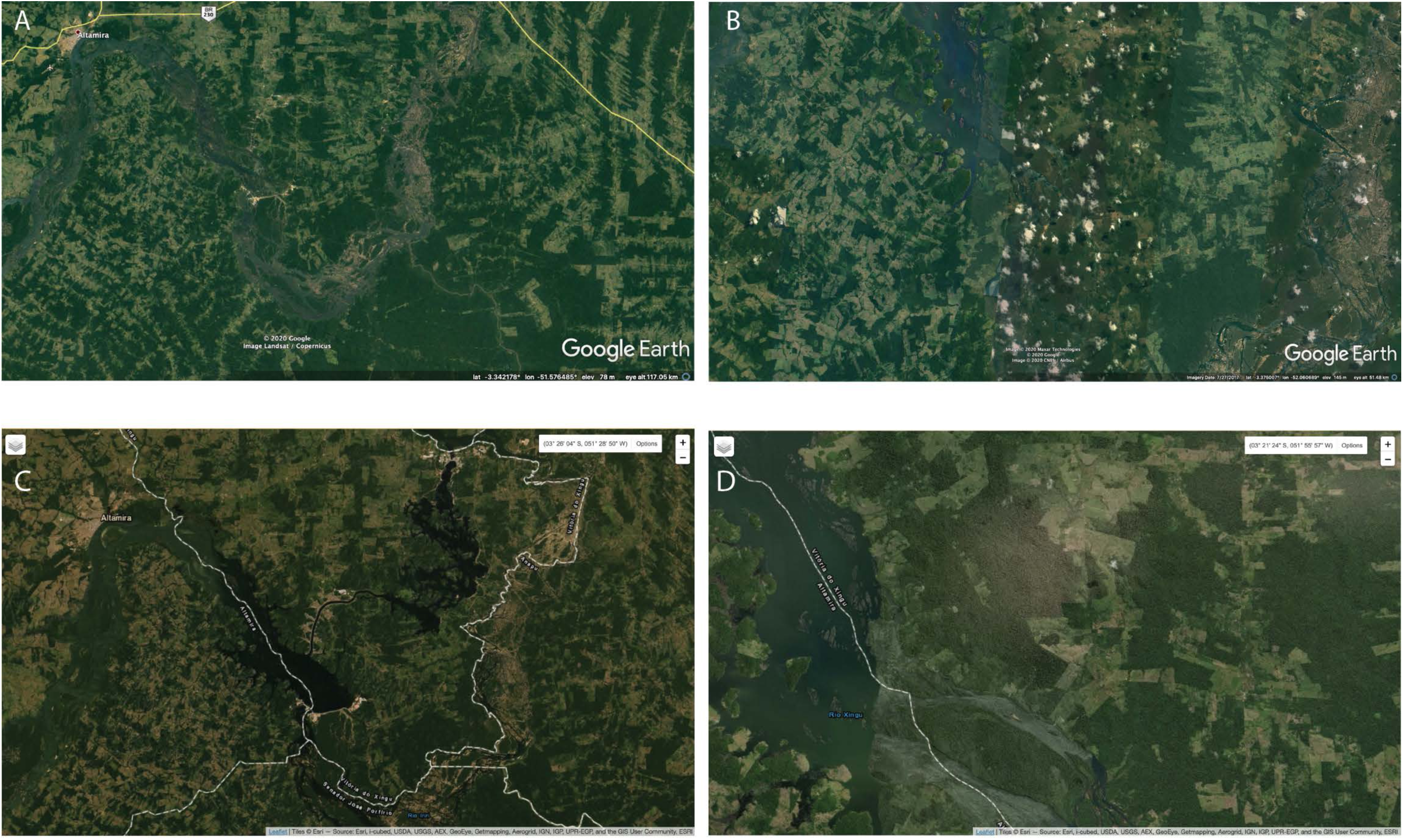
Example of the Google Earth representation accessed in 2020 of the Volta Grande at A) coarse scale and B) zoomed in to a finer level of detail. Given the status of the construction of the Pimental dam, the basemap image in A could not have been acquired later than 2014. In B, there is a split between imagery from 2017 showing the reservoir and imagery from 2010 with the original course of the river. C) Example of the basemaps used by the USGS EarthExplorer with imagery provided by ESRI at a coarse scale accurately showing the reservoirs from Belo Monte. D) zoomed in to a finer detail, the basemap representation from ESRI is also split in half with the reservoir only partially visible and the remainder of the area showing the original river’s course.

Despite its importance to all ecosystems and humanity, only ∼2.5% of the Earth’s water is freshwater, much of it stored in aquifers, glaciers and permanent snow cover (Stiassny 1996). Unsustainable usage is not only affecting the human population and global food security, but also the aquatic biota that depend upon those resources (Castello et al. 2013; Frederico et al. 2016). To compound the problem, terrestrial Land-Use and Land-Cover Change (LUCC) has a direct and grave impact on the aquatic ecosystems. But herein lies the problem: with few exceptions (e.g. Pekel et al. 2016), surface water is not the main focus in most global/regional/national classifications. And, unfortunately, the dedicated surface water classifications are rapidly out of date with large LUCC projects such as the Belo Monte dam not included. Users aware of the particular timeline of this hydroelectric project would know that the EC JRC/Google global surface water dataset for the 1984-2015 period is not appropriate for conservation questions past 2015. The inclusion of obsolete data into more recent land cover classifications (e.g. MapBiomas, TanDEM-X) can have grave ramifications. Similarly, basing conservation decisions on a modelled river network (e.g., HydroSHEDS) without first comparing the network to observed surface water may lead to incorrect assumptions about the ecosystems.

Land-Use and Land-Cover Change (LUCC) can have a dramatic effect on the release of terrestrial contaminants into freshwaters. For example, inorganic mercury is naturally found in Amazonian soils. However, modifications in Amazonian freshwater environments promote chemical reactions that result in its conversion to methylmercury (MeHg), the most toxic form of this metal (Berzas Nevado et al. 2010; Fearnside 2016b). Large construction projects, such as the Belo Monte dam, result in the decomposition of flooded terrestrial vegetation that deteriorates water quality and increases mercury’s bioavailability (Hsu-Kim et al. 2018). Former terrestrial islands in the segment of the river between Altamira and Pimental can be seen submerged following the flooding of the reservoir (Figure 11b,c). Given the gravity of the impacts of such large construction projects on entire regions, users must be careful in choosing the most appropriate datasets that accurately take into account their footprints.

As all datasets here demonstrate the proliferation of large-scale LUCC in the Xingu basin has continued since the 1980s. Urban centers (e.g., Altamira, São Felix do Xingu) have grown, but in parallel, the livelihoods of the population residing outside these primary centers have also undergone changes. These changes are a result of improved accessibility to markets alongside large scale expansion of the agriculture, energy, mining and forestry sectors within Brazil.

The Brazilian government’s new policies and talking points are a threat to conserving ecosystems across the country (Fearnside 2016a; Fernandes et al. 2017; Freitas et al. 2018; Tollefson 2018). For example, the current administration discredited the use of remote sensing by a national monitoring program (INPE: Instituto Nacional de Pesquisas Espaciais) to quantify forest loss in the Amazon (Escobar 2019b). With deforestation rates in Brazil continuing to increase at an alarming rate (Escobar 2020), our study shows that regardless of the variations in forest cover, all datasets concur that deforestation is still an issue in Brazil, and a serious threat to unique ecosystems like the Xingu river basin.

## ACKNOWLEDGEMENTS

This work was funded by the Natural Sciences and Engineering Research Council of Canada (NSERC) Discovery Grant Program to Kalacska and CNPq #486376/2013-3 and 309815/2017-7 to Sousa. PlanetScope and RapidEye imagery were provided through the Department of Geography at McGill University. We thank Márcio Vieira, Damilton Rodrigues de Costa (Dani) and Antônio Silva de Oliveira (Toninho) for their help in the field. We thank Dr. Mark Sabaj, Dr. John Lundberg and an anonymous reviewer for their comments to improve the manuscript.

